# Adaptation of endothelial cells to microenvironment topographical cues through lysyl oxidase like-2-mediated basement membrane scaffolding

**DOI:** 10.1101/2025.09.12.675854

**Authors:** Marion F Marchand, Noémie Brassard-Jollive, Claire Leclech, Jorge Barrasa-Fano, Yoann Atlas, Claudia Umana-Diaz, Apeksha Shapeti, Corinne Ardidie-Robouant, Tristan Piolot, Sabrina Martin, Philippe Mailly, Christophe Guilluy, Abdul I Bakarat, Catherine Monnot, Hans Van Oosterwyck, Stéphane Germain, Laurent Muller

**Author notes:** corresponding author: Laurent Muller : Center for Interdisciplinary Research in Biology (CIRB), College de France, CNRS UMR7241, INSERM U1050, 11 Place Marcelin Berthelot, Paris, France. contributed equally to this work (Noémie Brassard-Jollive, Claire Leclech). contributed equally to this work (Stéphane Germain, Laurent Muller). MFM: IBDM – CNRS UMR7288, Aix-Marseille Université, Campus de Luminy, Marseille, France NBJ: EFS, INSERM U1255, Strasbourg, France. YA: 3i Intelligent Imaging Innovations SAS, Paris, France. CUD: INSERM U944, CNRS UMR 7212, St. Louis Hospital Research Institute, Paris, France. CM: Centre de Recherche des Cordeliers, INSERM, Sorbonne Université, Université Paris Cité, Paris, France.

## Abstract

Basement membrane (BM) provides structural support and signaling platform for blood vessels. While its major structural components are required for vascular morphogenesis, integrating BM regulators, like the lysyl oxidase LOXL2, and BM assembly in cell response to microenvironement cues remain poorly understood. Here we study the early deposition and supramolecular assembly of BM components using correlative atomic force and fluorescence microscopy. The fibrillar deposition of fibronectin is gradually remodeled and associates with the collagen IV meshwork as it organizes into BM. We demonstrate that LOXL2 is deposited with both proteins and participates in their remodeling. Alteration of BM scaffolding by LOXL2-depletion affects focal adhesion maturation and cytoskeleton remodeling. This altered BM organization maintains stress fibers, affects the distribution and activation of mechanosensors and alters cell barrier properties. Furthermore, using 3D micro-printed substrates, we demonstrate that BM assembly regulates endothelial cell response to topographical constraint. We therefore propose a mechanism directly linking the scaffolding of BM components and adaptation to the topographical signals from the microenvironment.

## INTRODUCTION

Basement membranes (BM) are specialized, dense extracellular matrices (ECM) that outline tissues and play major roles in development and homeostasis(*1*, *2*). In addition to their structural support and signaling functions, recent studies implicate BM in the regulation of tissue morphogenesis through their mechanical properties(*3–5*). The vascular BM is a thin layer, less than 200 nm thick, composed of highly organized macromolecules that supports generation of the vascular wall at the interface between endothelial cells, pericytes and tissues(*6*). Endothelial cells initiate BM assembly as they invade the avascular microenvironment, and the loss of expression of major BM components, including laminin(*7*, *8*), collagen IV(*9*, *10*), and fibronectin(*11*, *12*), affects angiogenesis. Furthermore, altered expression or mutation of collagen IV and ECM remodeling enzymes affect vascular function and lead to the development of progressive vascular diseases such as small vessel disease(*13*), HANAC syndrome(*14*), vascular stiffening(*15*) and aneurysm(*16*, *17*). While vascular response to defective BM has been the subject of many studies, the integration of building the endothelial BM into cell response to microenvironmental stimuli, including topographical signals, remains to be understood.

Laminin is considered the building block of BM, thanks to its close interaction with the cell surface before connection to the collagen IV network, which self-assembles at the interface with the interstitial matrix(*18*), thereby producing a polarized structure(*19*). Numerous post-translational modification enzymes are involved in the processing of BM components during their synthesis as well as after their secretion, and 25% of the 160 human BM components identified are ECM regulators, including lysyl oxidase-like 2 (LOXL2)(*20*). LOXL2 is a secreted enzyme that catalyzes the deamination of hydroxylysines and lysines in collagen IV, thereby contributing to BM tensile force by cross-linking the 7S domains of collagen IV(*21*). LOXL2 also cross-links collagen I and elastin(*22*, *23*) and has been involved in matrix remodeling in cancer and fibrosis(*24*, *25*). During mouse retina vascularization and in zebrafish embryo, LOXL2 is expressed in endothelial tip cells and has been localized to the vascular basement membrane(*9*, *16*, *26*). We and others have shown that it regulates angiogenesis both in development(*9*, *27*) and in pathological contexts including cancer and post-ischemic revascularization(*9*, *28*, *29*). In the long term, LOXL2 is a susceptibility gene to aneurysm, including intracranial aneurysm(*17*, *30–32*). Our previous studies have shown that LOXL2 contributes to BM assembly without affecting the secretion of its components, and regulates capillary formation, without involving its enzymatic activity (*27*). However, the mechanisms connecting these two processes have yet to be elucidated.

In the present study, we first sought to characterize the early steps of the assembly of BM components and we report the remodeling of fibronectin deposited by endothelial cells into the collagen IV meshwork to initiate the vascular BM. We then characterized the involvement of LOXL2 in this process and analyzed how defects in BM assembly affect cell-ECM and cell-cell interactions. We show here that altering the remodeling of fibronectin and collagen IV after their deposition impacts the balance of cell-ECM and cell-cell adhesions as well as cell response to 3D printed micropatterns. We propose an original autocrine mechanism through which endothelial cells regulate their response to ECM topographical cues through the BM organization.

## RESULTS

### Early steps of endothelial BM assembly self-assembly

The vascular BM is generated during capillary sprouting, as illustrated at the vascular front of the postnatal mouse retina(*8*, *9*)(Fig. S1a-b). Fibronectin produced by astrocytes in the avascular microenvironment provides tracks for filopodia of endothelial tip cells, while higher amounts of fibronectin involved in *de novo* BM generation are detected at the surface of endothelial cells(*33*, *34*) (Fig. S1a). In contrast, collagen IV is restricted to endothelial cell surface, with tip cell protrusions projecting out of the collagen IV sheath (Fig. S1b). Similar sequential deposition of fibronectin and collagen IV was observed in a 3D capillary formation assay (Fig. S1c-d)(*35*). In order to acquire structural information to better understand the mechanisms involved in the early steps of the supramolecular assembly of these proteins, we here performed high-resolution analysis by correlative atomic force (AFM) and fluorescence microscopy. We have already characterized the polarized deposition of these proteins, together with laminin, perlecan and LOXL2, on the basal side of endothelial cells in culture(*27*). TIRF microscopy further demonstrated the direct polarized deposition of LOXL2 by these cells(*27*). Since the polarized deposition of matrix prevents direct access to the AFM, we first characterized the production of cell-derived matrix (CDM). A minimal culture time of 6 h after seeding was mandatory to achieve reproducible decellularization without alteration of the deposited material (Fig. S2a). To circumvent potential limitations of immunostaining, a direct assessment of LOXL2 deposition was performed using a LOXL2/GFP construct that colocalized with fibronectin and LOXL2 immunostaining (Fig. S2b and c, respectively). These experiments demonstrated that the generation of a homogeneous BM-like matrix required cell seeding at confluence and culture for a minimum period of three days (Fig. S2c). AFM topographic analysis of the organization of this BM-like matrix was thus performed between 6 h and 3 days after cell seeding. Deposition of nascent fibers measuring approximately 5 µm in length and 30 to 80 nm in height was detected at 6 h (Fig. 1a). These structures elongated and connected to form a uniform lattice by 24 h. Their diameter also increased 3-fold (Fig. 1b), suggesting the incorporation of additional proteins contributing to both fiber elongation and thickening. Forty-eight hours later, a homogeneous network 200 nm-thick had formed (Fig. 1a). The lateral extension of the fibrillar network continued throughout the process, covering up to 95% of the field at 72 h (Fig. 1b). While no significant change in overall height was observed between 6 and 24 h, major thickening of the ECM occurred between 24 and 72 h (Fig. 1b). BM deposition thus follows a biphasic growth pattern: an initial phase of deposition, elongation and interconnection of short fibrillar material, followed by a second phase of thickening through superimposition of fibrillar materials, ultimately forming a dense and cohesive network.

**Figure 1.**
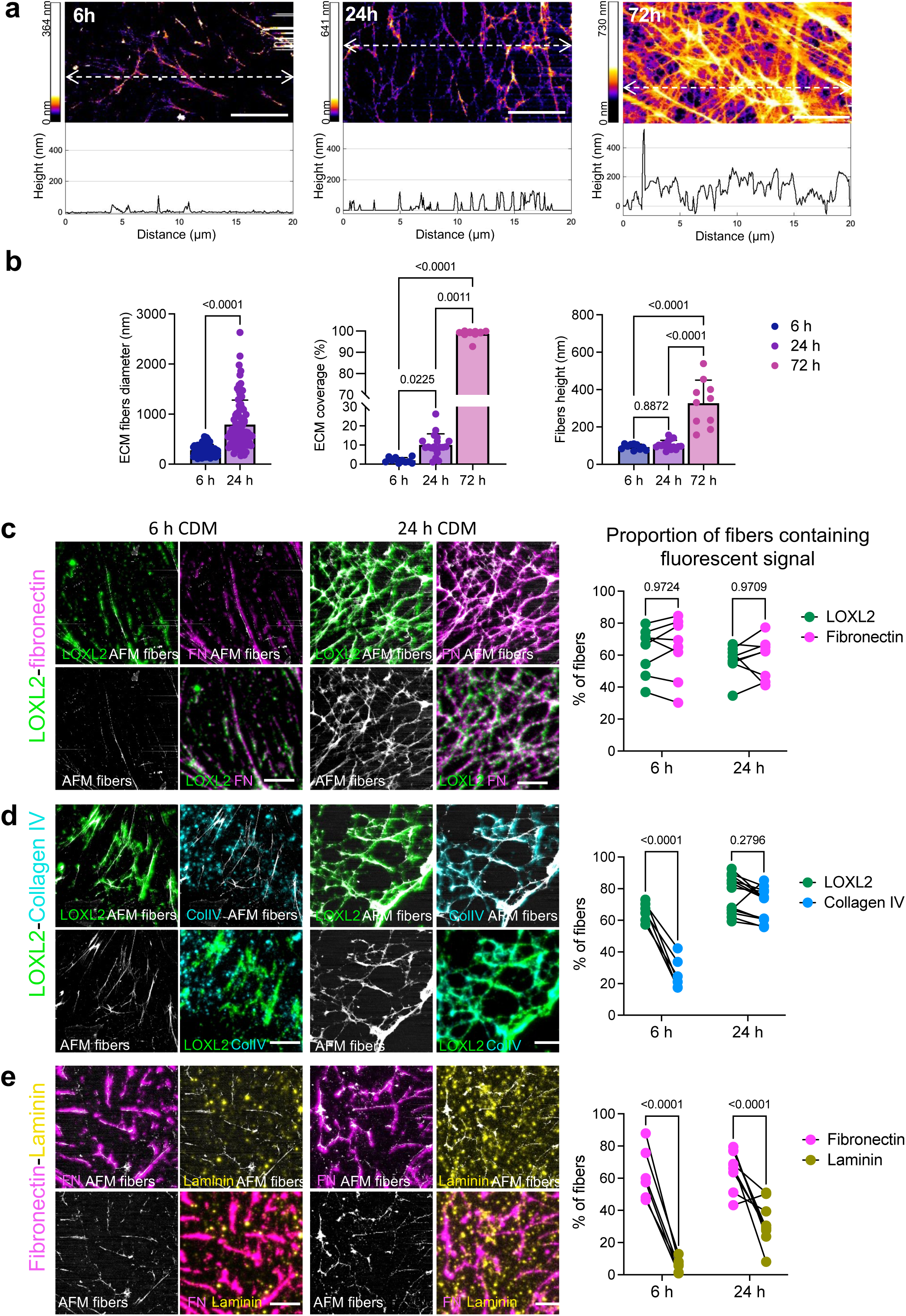
Endothelial basement membrane topography and assembly. **a:** Topographical analysis of basement membrane deposition using AFM imaging of CDM prepared 6, 24 or 72 h after seeding confluent endothelial cells. Height maps representative of at least 3 independent experiments are shown. White-dotted lines indicate positions of the scan lines of height. Scale bar: 5µm. **b:** ECM fiber diameter (µm), matrix coverage and mean fibers height (nm) were quantified in 2 (6 h) and 3 (24 and 72 h) independent experiments with a minimum of 4 maps/experiment. Mann-Whitney test for non-Gaussian distribution was performed for fiber diameters; Kruskal-Wallis test was performed for non-Gaussian distribution for matrix coverage; one-way ANOVA was performed for mean height. **c-e:** Topographical distribution of ECM components using correlative AFM-fluorescence analysis of CDM prepared from confluent cells cultured for 6 or 24 h and immunostained for LOXL2 (green) and fibronectin (magenta) **(c)**, or LOXL2 and collagen IV (cyan) **(d)** or fibronectin and laminin (yellow) **(e)**. Scale bar: 5µm. Quantification of the proportion of AFM-detected structures containing fluorescent signal was performed in 3 distinct experiments and 6 maps of 5x5 or 10x10 µm from each experiment. Statistical analysis was done using paired two-way ANOVAs.

The spatiotemporal distribution of BM components in these structures was analyzed by combining AFM topographic imaging with fluorescence microscopy (Fig. S3). We had previously shown the sequential co-localization of LOXL2 with fibronectin and collagen IV as well as their interaction in the matrix, using proximity ligation assay, and their direct binding using surface plasmon resonance(*27*). To decipher the mechanisms involved in BM organization, we here investigated CDM generated during 6 and 24 h of culture, as molecular crowding at longer time periods prevented reliable quantitative analysis. We first analyzed the distribution of fibronectin, collagen IV, laminin and LOXL2 immunostaining before performing AFM topography measurements and calculating the correlation rate (Fig. 1c-e). Fibronectin and LOXL2 were associated with ECM nascent fibrils as early as 6 h (64.2% ± 19% and 62.7% ± 15% respectively) and this association was stable over time as the microfibrillar network extended (Fig. 1c), which is consistent with LOXL2 being a component of the endothelial fibronectin-associated adhesome(*36*). While collagen IV was initially detected as a granular staining with no specific organization and in limited amounts in AFM-detected structures (26.1% ± 9%) at 6 h, the correlation increased over time to reach 70.4% ± 10% at 24 h (Fig. 1d). Throughout the process, laminin showed a diffuse distribution with very limited association with the structures detected by AFM despite a sharp increase in its deposition over time (Fig. 1e). We thus propose that fibronectin and collagen IV are initially deposited independently, with respectively high and low rates of association with LOXL2 in fibrillar structures detected by AFM. During the first 24 hours, BM network progressively organizes into interconnected supramolecular structures containing these three proteins. These findings provide insight into the early stages of BM organization, highlighting how the initial interactions between fibronectin, collagen IV and LOXL2 contribute to the formation of a dense network over time.

### LOXL2 drives BM spatio-temporal assembly

We then investigated the impact of LOXL2 depletion on the interactions between fibronectin and collagen IV during their early assembly, given that their mRNA expression and secretion are not deficient(*27*). After 6 hours, similar nascent fibronectin fibrils and granular collagen IV staining were deposited, regardless of LOXL2 expression (Fig. S4). However, the remodeling of these deposits into a common network at 24 h was impacted by LOXL2 depletion, as fibronectin was detected in fibrillar structures that were not co-localized with collagen IV (Fig. 2a)(*27*). Quantification of fibronectin motifs showed a higher alignment of fibronectin fibrils upon LOXL2 depletion without altering total length or fractal dimension of the fibrils (Fig. 2a). We then investigated the long-term consequences of this altered maturation by measuring the characteristics of CDM generated after 3 days. LOXL2 depletion had a high impact on ECM topography (Fig. 2b), replacing the homogeneous multi-layered network with large scattered aggregates interconnected by thin, sparse fibrillar structures. These matrix aggregates were more than twice the height of those deposited by control cells and contained both fibronectin and collagen IV (Fig. 2c-d). In contrast, the thin fibrillar material between these aggregates contained fibronectin but no collagen IV (Fig. 2c-d). Altogether, these data suggest that LOXL2 plays a crucial role in ECM maturation by promoting the association of fibronectin fibrils with the collagen IV network to form supramolecular complexes, thereby stabilizing these structures (Fig. 2e).

**Figure 2.**
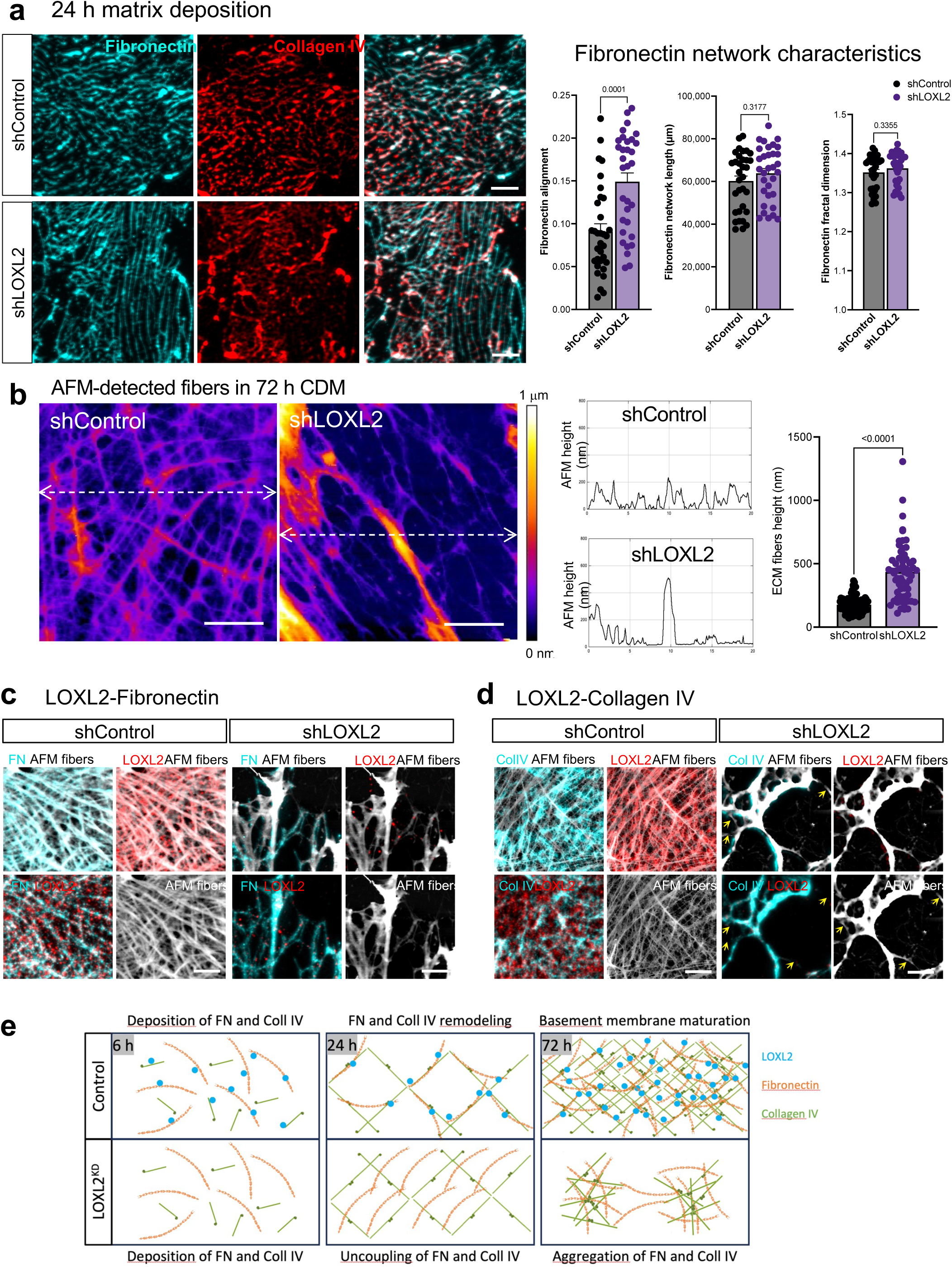
LOXL2 depletion affects spatio-temporal distribution of basement membrane components. **a:** LOXL2 is required for fibronectin remodeling at 24 h. Immunostaining of fibronectin (cyan) and collagen IV (red) in CDM generated by control or LOXL2-depleted endothelial cells over 24 h. Scale bar: 10 µm. Alignment, total length and fractal dimension were quantified for fibronectin staining at 24 h in n = 32 images per condition, from 3 independent experiments. Statistical analysis was performed using Mann-Whitney tests. **b:** Topographical AFM analysis of CDM generated by control or LOXL2-depleted endothelial cells over a period of 72 h. White dotted line indicates position of the scan lines of height shown on the right. Scale bar: 5 µm. Height of individual fibers (nm) was extracted from scan-lines in 2-5 images from each of 4 distinct experiments (total of 60-100 fibers) and unpaired Student t-test was performed. Data is represented as mean + SD. **c-d:** LOXL2-depletion affects overall basement membrane organization at 72 h. Correlative AFM-fluorescence analysis of CDM generated by control or LOXL2-depleted endothelial cells over 72 h. CDM were immunostained for fibronectin **(c)** or collagen IV **(d)**. Yellow arrows point structures detected by AFM and lacking collagen IV staining. Scale bar: 5µm. **e:** Schematic representation of basement membrane assembly by cells expressing (top) or not (bottom) LOXL2 at 6, 24 and 72 h.

### BM regulates cell-cell and cell-ECM interactions through an autocrine mechanism

The strong impact of LOXL2 depletion on fibronectin and collagen IV organization between 24 and 72 hours prompted us to investigate how the altered ECM scaffolding affects cytoskeleton remodeling and cell shape. F-actin was remodeled in response to LOXL2 expression (Fig. 3a). In control cells, the actin cytoskeleton was predominantly subcortical, distributed at the vicinity of cell-cell junctions. In contrast, LOXL2-depleted cells showed stress fibers bundles oriented along the cell axis. We evaluated the orientation of F-actin and observed a preferential alignment in these cells (Fig. 3a). While control cells acquired a typical cobblestone shape and organization as detected with VE-Cadherin distribution, LOXL2-depleted cells underwent major shape modifications resulting in aligned and elongated cells with 30% reduction in circularity (Fig. 3b). Cell shape modification was associated with remodeling of cell-cell junctions. The morphology of adherens junctions was classified into three categories defined as straight, reticular and serrated structures (Fig. 3c). In line with the changes observed in cell morphology and the associated contractile cytoskeleton, the proportion of straight junctions increased 1.6-fold in cells with altered BM (Fig. 3c). Consistent with these observations, the barrier properties measured 72 hours after seeding were also altered in LOXL2-depleted cells (Fig. 3d).

**Figure 3.**
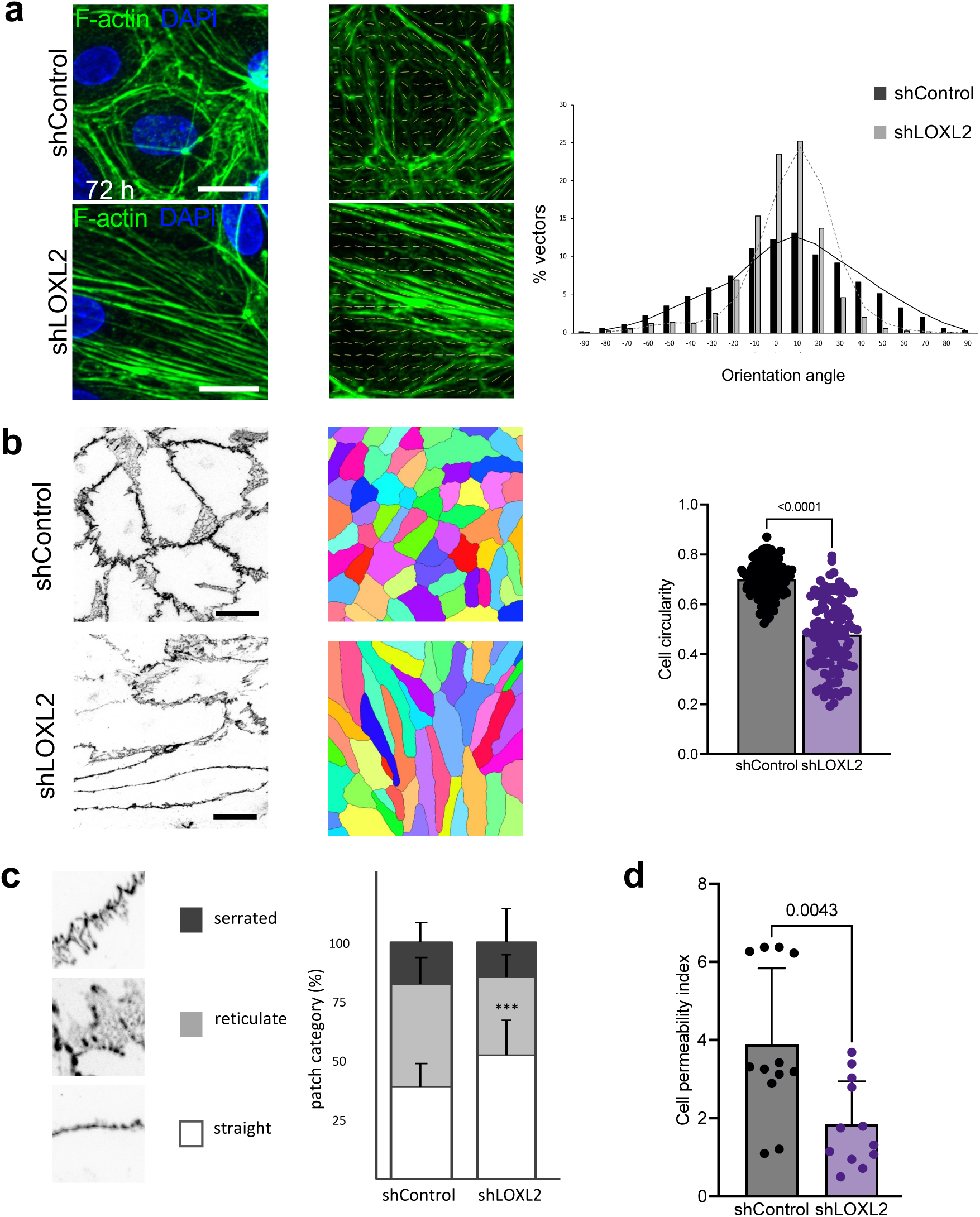
LOXL2-mediated basement membrane assembly drives cytoskeleton organization and cell-cell junctions. **a:** Detection of F-actin with phalloidin-Alexa Fluor 488 was performed at 72 h. Scale bar: 20 µm. Cytoskeleton orientation was assessed in 3 independent experiments (n = 10 fields of view of 60x60 µm per condition). **B:** Immunostaining of b-catenin 72 h after seeding control or LOXL2-depleted confluent endothelial cells. Cell circularity was calculated in n = 144 (shControl) and n = 122 (shLOXL2) cells, from two distinct experiments. Statistical analysis was done using unpaired t-tests. **c:** Evaluation of cell-cell junction morphology was performed on VE-Cadherin-stained endothelial monolayers after sorting in straight, reticulate and serrated categories. Data are mean ±SD of 3 independent experiments each producing 4-7 fields of view per condition. *p* values reported were calculated for straight junctions using unpaired t-tests between control and LOXL2-depleted cells in control conditions (*** p < 0.0005). **d:** Barrier properties of endothelial monolayers were measured after 72 h in culture. Data are means of 4 experiments performed in duplicate or quadruplicate culture wells (n = 12). Statistical analysis was done using unpaired t-test.

Given the known cross-talks between cell-ECM and cell-cell adhesions in endothelial cells, we investigated the distribution of β1 integrin, the fibronectin (α5β1) and collagen (α1β1 and α2β1) receptor, and associated mechanosensors. After 24 hours, β1 integrin was co-localized with fibronectin fibrils and its distribution did not depend on LOXL2 expression (Fig. 4a). The sharp increase in fibronectin network density over time was accompanied by a redistribution of adhesions containing β1 integrin (Fig. 4a). While the redistribution of β1 integrin over time corresponded to the organization of actin described above (Fig. 3), it did not reflect the remodeling of the fibronectin network. Upon LOXL2 depletion, β1 integrin was maintained in fibrillar adhesions co-localized with the sparse thin fibers of fibronectin described above (Fig. 2c and d). Interestingly, β1 integrin was virtually absent from the large fibronectin aggregates (Fig. 4a).

**Figure 4.**
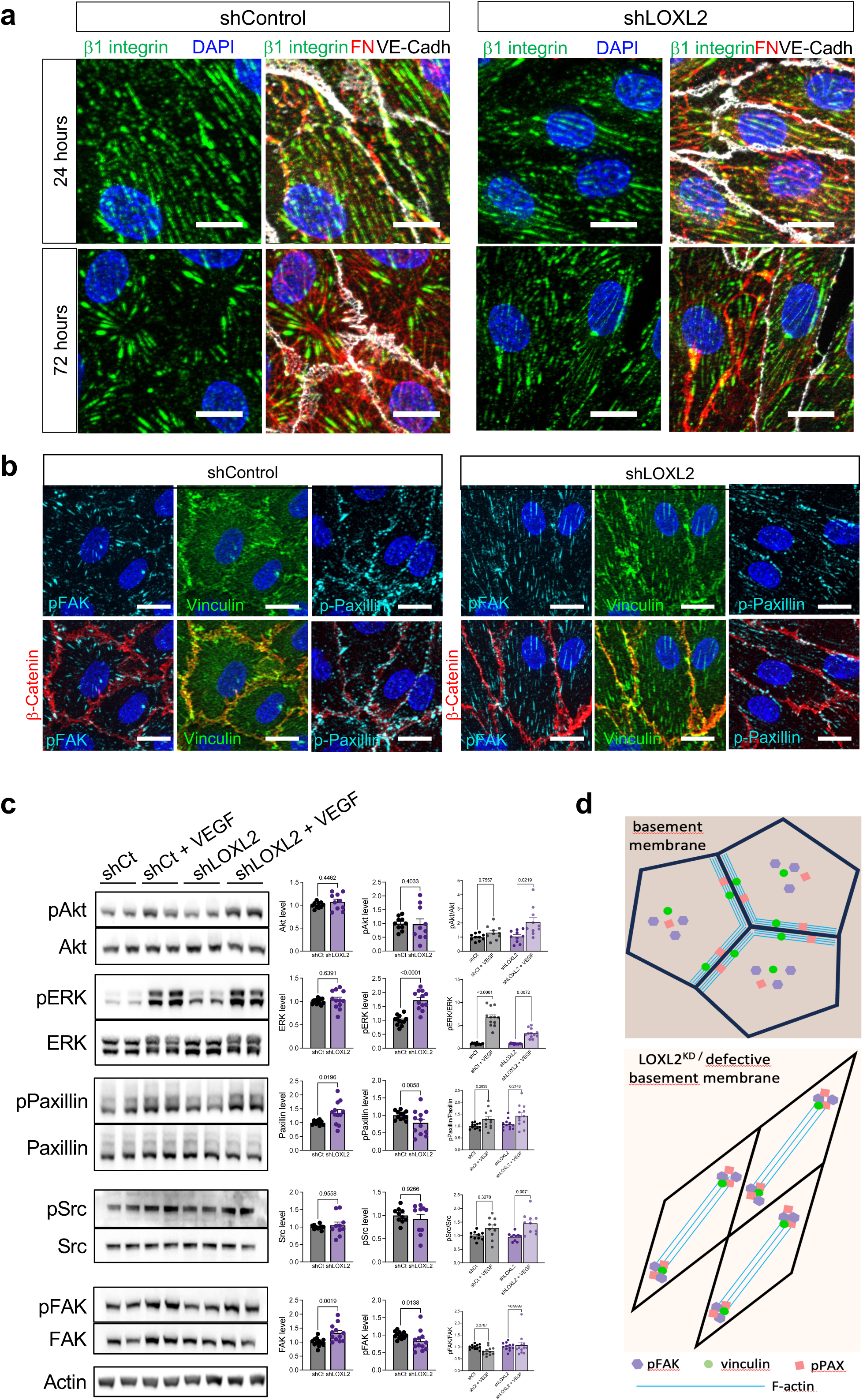
Defective basement membrane scaffolding alters maturation of cell-ECM adhesions and cell-cell junctions. **a:** Immunostaining of b1 integrin (green), fibronectin (red) and VE-cadherin (white) in control or LOXL2- depleted endothelial cells 24 or 72 h after seeding confluent cells. **b:** Immunostaining of pY397-FAK (pFAK – cyan), vinculin (green), pY118-paxillin (pPAX – cyan) and b-catenin (red) in control or LOXL2- depleted endothelial cells 72 h after seeding confluent cells. Nuclei were detected with DAPI (blue). Scale bar: 10 µm. **c:** Control (shCt) or LOXL2-depleted (shLOXL2) endothelial cells were cultured 72 h before cell lysis and western blotting for the indicated proteins. The level of expression (left column), basal phosphorylation (center column) and VEGF-induced phosphorylation (right column) were quantified in 4 independent experiments performed in duplicate culture wells (n = 8). Statistical analysis was done using Kruskall-Wallis tests. **d:** Schematic of cytoskeleton remodeling and distribution of adhesion proteins in cell-ECM and cell-cell junctions in control and LOXL2-depleted cells.

In order to rule out the possibility that LOXL2 depletion intrinsically alters the mechanotransduction properties of endothelial cells, we assessed their response to controlled ECM stimuli in the absence of ECM deposition. Adhesion was analyzed within one hour after seeding cells on fibronectin or collagen I coated surfaces. The overall architecture of the actin cytoskeleton and cell spreading were similar in control and LOXL2-depleted cells (Fig. S5a). Paxillin staining was similarly distributed in nascent adhesions at the cell periphery and matured into focal adhesions towards the cell center. The size and morphology of these adhesions were comparable between conditions, indicating that LOXL2 depletion did not affect focal adhesion formation or structure (Fig. S5a). Similar observations were made for vinculin (Fig. S5b). To further demonstrate that altered BM generated by LOXL2-depleted cells was responsible for the adhesion defaults described above, control cells were plated on control or LOXL2-depleted CDM. In control conditions, the distribution of paxillin and pY397-FAK was consistent with previously described patterns on fibronectin or collagen I surface coating (Fig. S5a and c). On the other hand, matrices prepared from LOXL2-depleted cells did not promote maturation of nascent adhesions into focal adhesions in control cells, even though they supported cell adhesion and spreading (Fig. S5c).

Distribution and activation of mechanosensors were thus investigated after 3 days of BM generation (Fig. 4b-c). While pY397-FAK was detected in focal adhesions of control cells with an isotropic organization, we observed elongated aligned adhesions in LOXL2-depleted cells (Fig. 4b), corresponding to β1 integrin distribution (Fig. 4a) and cytoskeleton remodeling (Fig. 3c). While vinculin was predominantly co-localized with β-catenin at cell-cell junctions as expected in control cells, its staining was also maintained at focal adhesions on LOXL2-depleted matrices (Fig. 4b). Phospho-Y118-paxillin was localized in the vicinity of β-catenin staining at the cell-cell junctions of control cells (Fig. 4b). This distribution was altered in LOXL2-depleted cells with persistent staining of focal adhesions as observed with vinculin. The presence of vinculin, activated FAK and paxillin in elongated cell-ECM adhesions in LOXL2-depleted cells led us to further investigate integrin signaling in these culture conditions (Fig. 4c). FAK and paxillin expression were increased in response to LOLX2 depletion, while their basal phosphorylation was reduced. On the other hand, LOXL2 down-regulation did not impact the expression of Akt, ERK, and Src (Fig. 4c), but ERK basal phosphorylation was increased by 1.6-fold, in agreement with the cytoskeleton contractile phenotype observed in these cells. We also investigated cell response to the angiogenic growth factor VEGF. Whereas VEGF-mediated phosphorylation of ERK was lower in cells depleted for LOXL2, the total level of activated ERK was maintained in these cells as a result from higher basal activation. In addition, Src and Akt activation was increased in LOXL2-depleted cells, indicating higher activation of the proliferation/cell survival signaling in these cells. In addition, the redistribution of mechanosensors and increased activation of ERK and Src support the modulation of endothelial barrier properties described above (Fig. 3c-d).

Together, our data thus link BM remodeling to maturation of focal adhesions, cytoskeleton organization and cell morphology. They more particularly suggest that BM assembly regulates the balance of mechanosensors at focal adhesions and adherens junctions (Fig. 4d), with functional consequences on junction organization and barrier properties.

### BM assembly mediates 3D cell organization

To investigate how altered organization of the BM affected cell response to 3D microenvironment., we first analyzed cell behavior in response to topographical cues. Endothelial cells were seeded on PDMS micro-grooved substrates featuring grooves 2 µm wide and spaced 2 µm apart, with a depth of 1 µm. These dimensions correspond to the fibrillar structures that were detected by AFM at 6 h, as well as to the organization of matrix in tissues and hydrogels(*37*, *38*) (Fig. 5a and S6a). Control cells initially organized in a confluent monolayer aligned along the microgrooves at 24 hours and gradually adopted an isotropic orientation without any significant change in cell circularity over the following 48 hours (Fig. 5b-c). This was accompanied by reorganization of the actin cytoskeleton, shifting from bundles of stress fibers aligned along the cell axis to cortical actin associated with intercellular junctions, and by the similar redistribution of β1 integrin as observed on non-patterned culture substrate (Fig. 5b and d). In contrast, LOXL2-depleted cells aligned more strictly with the grooves at 24 h and maintained this orientation and elongated shape over the following 48 h, indicating a sustained response to topographical cues. Consistent with this, stress fibers and elongated focal adhesions containing β1 integrin were preserved in LOXL2-depleted cells (Fig. 5b and d). Similar cell behavior was observed on 5µm ridges and grooves that were also only 1µm deep (Fig. S6). Control cell behavior and focal adhesion remodeling were associated with an isotropic organization of fibronectin and collagen IV extending over the groove pattern, whereas both proteins were detected as sparse aggregates in the case of LOXL2-depleted cells (Fig. 5e and S6f). These results indicate that confluent endothelial cells initially align along microgrooves consistent with constrained contact guidance, until BM deposition releases them from these topographical cues. We then assessed cell behavior in a model of capillary formation in 3D collagen hydrogel(*33*). Twenty-four hours after seeding, endothelial cells have stacked up fibrillar collagen I at their surface and initiated deposition of patches of collagen IV at their surface (Fig. S7a). Cell spreading and compaction of collagen I at the surface of endothelial cells remained unchanged when LOXL2 was depleted (Fig. 6b), suggesting that this enzyme does not directly impact the mechanical properties of the cells. Consistently, traction force microscopy experiments demonstrated that cell spreading and average traction forces both increased with stiffness of the culture support, but that these features were not affected by the level of expression of LOXL2 (Fig. S8a). Cell response to stiffness was also assessed by performing AFM stiffness-clamp experiments (Fig. S8b). This setup enables the evaluation of mechanosensing by tuning the apparent stiffness of the cell microenvironment in real time(*39*). Force rate was recorded dynamically while applying a broad range of stiffness values, ranging from 10mN/m to 400mN/m during a continuous single-cell traction force experiment. Similar increasing traction rates were measured as the applied stiffness increased, regardless of whether cells expressed LOXL2 (Fig. S8b). Altogether, these results indicate that LOXL2 depletion does not affect the early stages of cell adhesion and spreading, nor does it affect cell mechanosensing or contractility. Seventy-two hours after seeding in collagen hydrogels, cells have self-organized into capillary-like structures and deposited collagen IV in the BM at the interface with relaxed collagen I (Fig. S7b), suggesting a shift of cell interactions with hydrogel collagen I to their own BM. LOXL2-depleted cells exhibited impaired organization characterized by reduced diameter and fewer connections when BM generation was impaired (Fig. 6c) as previously demonstrated(*9*, *27*). While control cells deposited collagen IV in the BM, with lower amount at the surface of the leading cell, LOXL2 knock-down was associated with poor deposition of collagen IV and by the presence of empty sleeves of faint and discontinuous collagen IV staining (Fig. 6d). These defects were accompanied by a significant increase in cell death, reaching up to 70% for LOXL2-depleted cells (Fig. 1a and d). However, such elevated cell fate was specific to the 3D culture context, as no difference in cell death was measured when cells were cultured on 2D tissue culture plastic or 2D collagen I coated surfaces (Fig. S9). Taken together, these data demonstrate that BM deposition by endothelial cells alleviates their dependance on external topographical cues, instead providing them with a self-generated 3D microenvironment that enables cell self-organization into capillary-like structures.

**Figure 5.**
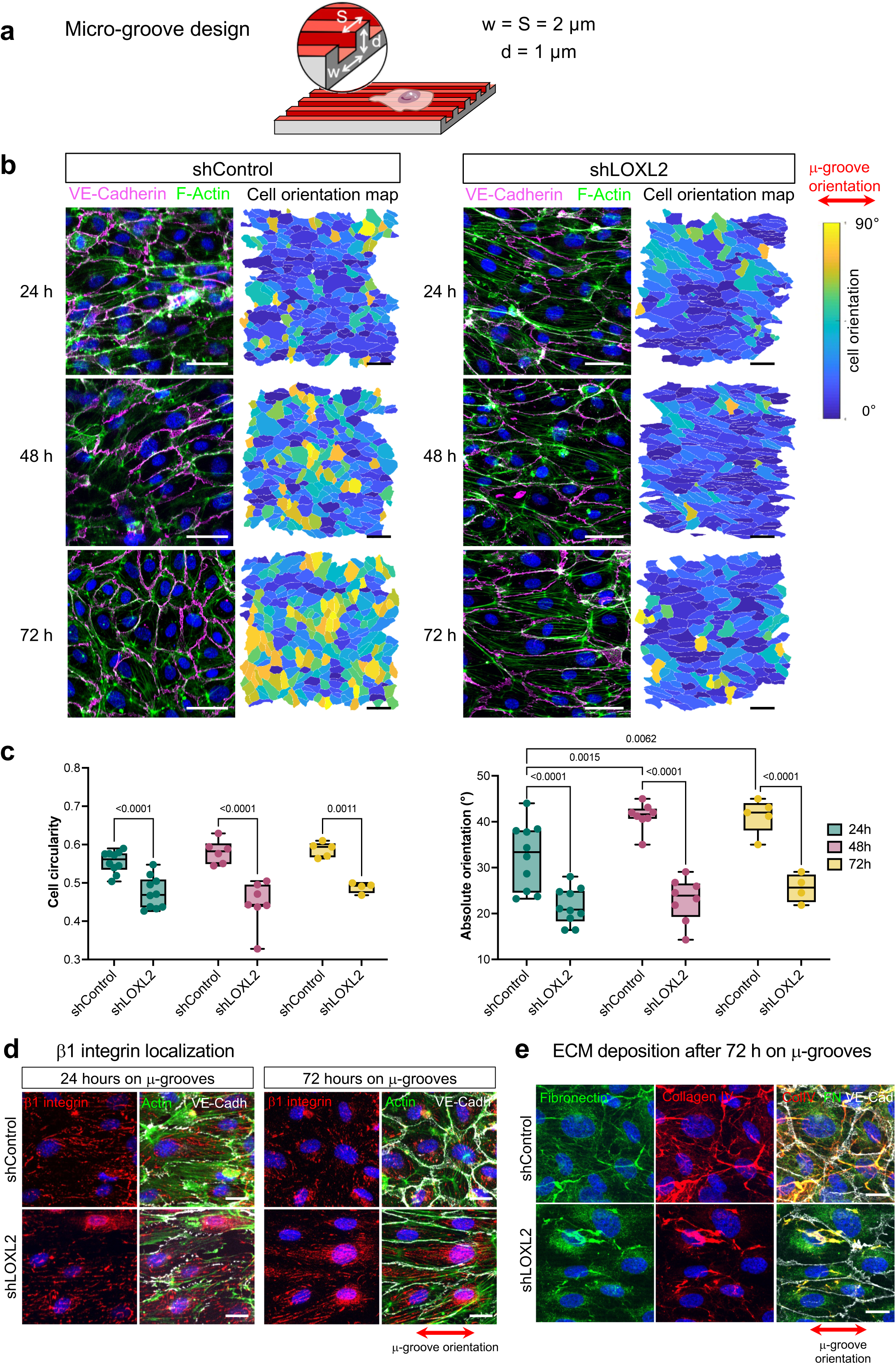
Defective basement membrane scaffolding affects cell response to topography. **a:** Schematic of the microgrooved culture substrate. Two µm wide (w) and 1 µm deep (d) grooves were spaced (S) by 2 µm. **b:** Immunostaining of VE-cadherin (magenta) and staining of F-actin with phalloidin-Alexa Fluor488 (green) 24, 48 or 72 h after seeding control or LOXL2-depleted confluent endothelial cells on microgrooved substrates. Scale bar: 50 µm. Cell orientation maps are shown for each condition. Scale bar: 100 µm. **c:** Cell shape and orientation were measured after segmentation and image analysis of VE-cadherin staining in 3 independent experiments. **d:** Immunostaining of b1 integrin (red) and VE-cadherin (white) in cells cultured for 24 and 72 h on micro-grooved substrates. F-actin was stained with phalloidin-Alexa fluor488 (green). Scale bar: 20 µm. **e:** Immunostaining of fibronectin (green), collagen IV (red) and VE-cadherin (white) in cells cultured for 72 h on micro-grooved substrate. Nuclei were detected with DAPI (blue). Scale bar: 20 µm. Red double arrows indicate the orientation of the microgrooves.

**Figure 6.**
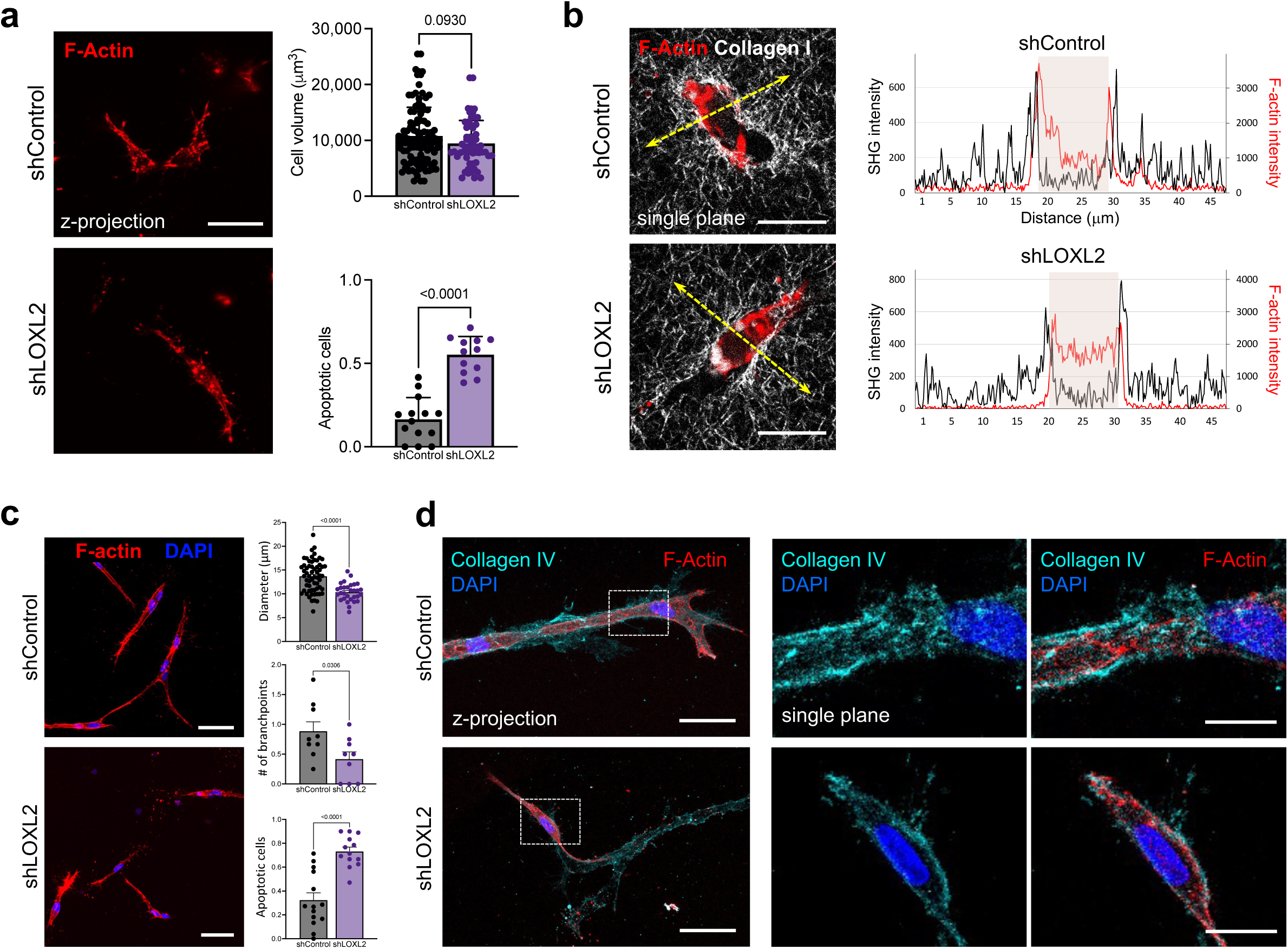
Basement membrane deposition and 3D vascular morphogenesis. **a-b:** Endothelial cells were cultured in 3D collagen I hydrogels for 24 h before F-actin staining with phalloidin (red). Scale bar: 50 µm. Cell volume (µm^2^) was calculated in n = 59-111 cells/condition. Proportion of dead cells was assessed by counting fragmented nuclei over total cell nuclei in n = 12-14 images per condition. All parameters were quantified in 3 independent experiments each performed in duplicate hydrogels. Statistical analysis was performed using unpaired t-tests and Mann-Whitney test. Collagen I was visualized using SHG in 2-P microscopy (white-right panel) **(b)**. Scan lines of intensity are shown to illustrate collagen I compaction at the cell surface. Scale bars: 10 µm. **c-d:** Endothelial cells were cultured in 3D collagen I hydrogels for 72 h before F-actin staining with phalloidin (red). Scale bars: 50 µm. Diameters and number of connections of capillary-like structures were calculated in n = 30-65 cells per condition. Number of dead cells was assessed by counting fragmented nuclei in approximately 15 fields of view/condition. All parameters were quantified in 3 independent experiments each performed in duplicate hydrogels. Statistical analysis was performed using unpaired t-tests. Collagen IV (cyan) was detected using confocal imaging and Z-stack projections are displayed (left panel) **(d)**. Squared box corresponds to the zoomed area displayed as single z plane in the right panel. Scale bars: 50 and 10 µm, respectively. For all graphs, data is represented as mean + SD and p-values are indicated.

## DISCUSSION

We here present a detailed analysis of the early steps of fibronectin and collagen IV scaffolding in the BM with a focus on the involvement of LOXL2. These proteins are first deposited in independent fibrillar and granular structures, respectively. Collagen IV is then remodeled into a network that incorporates fibronectin but does not resemble any of these initial deposits. The collagen IV-driven reorganization of fibronectin has previously been proposed(*40*, *41*). Our observations are also consistent with the fibronectin localization at the surface of tip cells extending protrusions out of the collagen IV sheath assembled by stalk cells during retina vascularization(*8*, *9*). Most remarkably, the colocalization of fibronectin to self-assembled collagen IV structures is regulated by LOXL2. In contrast, LOXL2 does not affect laminin deposition, confirming our previous observations(*27*). Since BM is a polarized structure with laminin associated with the surface of endothelial cells (*18*, *19*), we propose that LOXL2 affects BM organization at the interface between collagen IV and the interstitial matrix. Our findings will thus be impacted by the interactions of the newly formed basement membrane with the matrix in the newly vascularized microenvironment. In line with these data, LOXL2 and collagen IV α1 and α2 are both components of the fibronectin-bound endothelial adhesome, from which laminin α4, ß1 and ψ1 are excluded(*36*). Although the association of fibronectin with collagen IV has been described for long(*42*, *43*), no binding site has yet been identified for their direct interaction. Our findings suggest that LOXL2 could participate in scaffolding both proteins, either directly as we have detected its binding to each protein using surface plasmon resonance, or indirectly through another BM component, as we could not identify a three protein complex (*27*).

Maturation of BM is further achieved by accumulation of multiple layers of material resulting in a thicker and denser network. Multicellular mechanisms involving migrating cells have already been described for the generation of BM during egg chamber elongation in drosophila(*44*, *45*) and neural crest cells migration(*46*). Consistently, we have already described the persistent exocytosis of LOXL2 and its incorporation in preexisting matrix by migrating endothelial cells(*27*), which fits with endothelial cell migration during sprout elongation. Sequential deposition and remodeling of fibronectin and collagen IV is further supported by the deletion of β1 integrin in endothelial cells, which prevents deposition of fibronectin but not of collagen IV, yet later results in loss of endothelial BM (*47*). The β1 integrin-mediated formation of fibronectin fibrils requires its synthesis by endothelial cells(*11*, *48*) and is essential as a BM organizer(*43*, *47*). We observed decreased co-localization of β1 integrin with fibronectin as BM matures, suggesting that increasing association of fibronectin with LOXL2 and collagen IV could shield fibronectin from β1 integrin-mediated cell tractions. Such process has been described in interstitial ECM, where fibronectin interactions with fibrillar collagen are regulated by the tension level of fibronectin and the associated cellular traction forces mediated by integrin binding(*49*). In agreement with this hypothesis, blood vessels are indeed lined with untensed fibronectin(*50*). In the absence of LOXL2 and the associated stabilization of collagen IV-fibronectin association, both proteins accumulate in large and disorganized aggregates, with thin fibrillar structures containing fibronectin only. Consistently, defective BM production alters endothelial cell morphology and cytoskeletal remodeling, even though the cellular properties of adhesion and mechanotransduction *per se* are not affected. It is thus the defective BM that affects activation and distribution of mechanosensors including vinculin and paxillin, which are retained at cell-matrix elongated adhesions in addition to their distribution at the cell-cell junctions(*51–54*).

Lower level of activation of paxillin supports stabilization and decreased turnover of focal adhesions(*55*), and regulates the balance between adhesions and junctions through interactions with FAK or β-catenin, respectively(*56*). As a result, the shift in the balance of mechanosensors between adhesions and junctions affected endothelial barrier properties, which could affect vascular function in the long term (*57*). Local geometric control of ECM and ECM ligand micropatterning have been known to drive cytoskeleton organization and cell fate for long(*58*, *59*). However, the integration of building up the BM as a response to microenvironment cues is still a key factor to characterize(*41*), considering the importance of topography for patterning microvascularization. When seeded in 3D gels, cells first spread by exerting traction forces on collagen fibers. Impeding BM deposition at the interface with collagen I prevents further capillary morphogenesis over the next days and results in high levels of cell death. Collagen IV depletion also prevented capillary formation in 3D hydrogels(*9*). This process however does not occur in 2D culture, including on collagen I coated surface, unless cell shape and spreading is decreased and thus triggers apoptosis of endothelial cells(*58*). Cell-mediated building up of the BM thus participates in the regulation of cell fate. We also demonstrate that this process enables cells to overcome topographical patterns. While confluent endothelial cells seeded on 3D anisotropic substrates first align in a similar manner to single cells(*60*, *61*), we have shown that generation of BM over the following days allows cells to overcome these topographic constraints (*62*). In contrast, vascular smooth muscle cells produce a fibrillar extracellular matrix organized along the grooves(*63*). Given that secretion of BM components is not altered(*27*), our findings demonstrate that the autocrine BM organization is crucial for cells to overcome topographical constraints.

Taken together, our data thus demonstrate that BM generation is a key event mediating cell 3D organization in response to microenvironmental cues. Such mechanism participates in the cell capacity to organize into capillary through the shift of interactions with the interstitial ECM to interactions with vascular BM. Furthermore, the dysregulation of adherens junctions and barrier properties associated to LOXL2 mediated defaults in BM supports its involvement in the progression of long-term microvascular BM diseases. LOXL2 has indeed been identified as a susceptibility gene to aneurysm, and more specifically intracranial aneurysms(*17*, *30*, *31*, *59*), and was associated to BM defaults including collagen IV deposition in the progression of retina microaneurysms(*16*).

## METHODS

### Antibodies

Primary and secondary antibodies are listed in table 1.

**Table 1:**
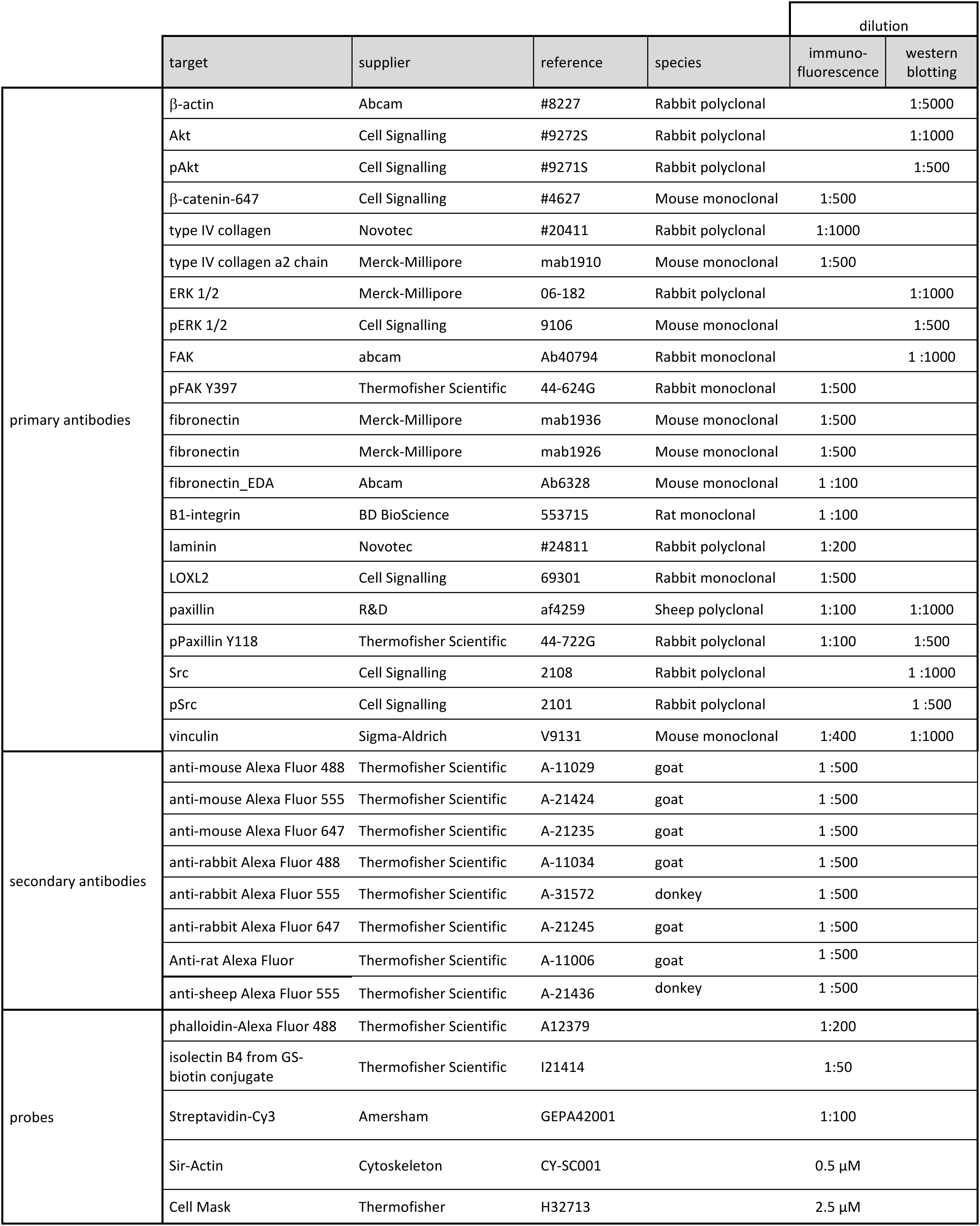
Antibodies and probes.

### Cell culture and lentiviral transduction

Human umbilical vein endothelial cells (HUVECs) were prepared and grown as already described(*35*) in endothelial cell growth medium (ECGM2, Promocell) and on collagen I-coated substrates. Cells expressing LOXL2/GFP or down-regulated for LOXL2 have previously been described(*27*). Experiments were performed using HUVECs between passages 2 and 5. For coating experiments, collagen I (Corning) or human plasma fibronectin (Sigma-Aldrich) were respectively diluted in 20 mM acetic acid or PBS before coating culture dishes for 1h at 37°C.

### 3D collagen hydrogels

Endothelial cells were seeded in 2 mg/mL collagen I extracted from rat tail as previously described(*35*). Hydrogels were cast in ibidi µ-inserts and cultured from 1 to 3 days in fibroblast conditioned medium(*35*). 3D samples were fixed with 4% PFA for 30 min and incubated overnight with primary antibodies or phalloidin coupled to complementary Alexa Fluor (1:200; ThermoFisher Scientific). Images were acquired with a 20X/0.8-WD 550 1m or 40X/1.1-WD 620 1m objective mounted on a CSU-W1 spinning-disk confocal head (Yokogawa) mounted on an inverted Z1 Axio Observer Zeiss microscope. Images were segmented using a 3D mask (Imaris, Bitplane) and further analyzed with a Matlab (MathWorks) in-house software (named 3D-Skel)(*35*). For imaging of endothelial cell organization and collagen remodeling, we used two-photon (2-P) microscopy on a Leica DMI6000 Upright SP5 confocal microscope using a 25X/0.95-WD 2.5 mm water immersion objective (Leica). Two-photon excitation was performed using a Mai Tai SP laser (Spectra Physics). Second-harmonic generation (SHG) signal was collected by external non-descanned detectors at 440 nm, using 2-P excitation at 880 nm.

### Preparation of cell-derived extracellular matrices

Endothelial cells were seeded at confluency in low IbiTreat Petri dishes (Ibidi) and cultured in ECGM2 for the indicated amounts of time. All steps were performed on ice using ice-cold solutions. Cells were washed with PBS and detached using a solution containing 0.3% Triton X-100 and 20 mM ammonium hydroxide (Sigma-Aldrich) in PBS. CDM were washed three to four times with PBS before further processing. Samples were checked under a light microscope to ensure proper decellularization.

### Immunofluorescence and image analysis

Endothelial cells were seeded in 8-well ibiTreat µ-slides (Ibidi) or in low ibiTreat Petri dishes (Ibidi) and cultured for 6 to 72 h before fixation with 4% paraformaldehyde and permeabilization with 0.5 % Triton-X-100. Appropriate primary and secondary antibodies were incubated in the presence of 1% normal goat serum and 0.01% Triton-X-100. Z-stacks were acquired with either a CSU-W1 spinning-disk confocal (Yokogawa) on an inverted Z1 Axio Observer microscope or a Z1 Axio Observer epifluorescence microscope equipped with an Apotome2 (Zeiss) using 63x-NA1.4 or 40x-NA1.1 objectives. Due to the lack of evenness of the culture supports, projections were analyzed using Image J software. Fibronectin matrix metrics were obtained using the TWOMBLI imageJ pipeline(*64*). Cell circularity was calculated using the MorphoLib plugin. Analysis of actin cytoskeleton orientation was performed on 50x50 µm cropped images centered around cell nuclei, using the OrientationJ plugin to generate vector fields. Computation of the data included normalization of vector distribution after subtraction of the median value in each image. Focal adhesion metrics were quantified using an in-house macro based on the Analyze Particles function of image J. Analysis of junction morphology was performed manually using a method adapted from Neto and collaborators(*65*). Five to seven 211x211 µm^2^ fields of view were acquired for each condition and 100 patches of 10x10 µm^2^ were randomly extracted from each image using in house script adapted from Neto and collaborators. Morphology of the cell-cell junctions in each patch was manually sorted into three categories corresponding to straight, reticulate or finger/serrated junctions.

### Traction force microscopy

Endothelial cells were seeded on top of fibronectin-coated home-made polyacrylamide hydrogels of various stiffnesses containing fluorescent beads, as already described(*66*, *67*). Briefly, gels were cast in APTS- and glutaraldehyde-treated (Sigma-Aldrich) 35 mm glass-bottom Petri dishes (MatTek). Hydrogels were produced using 625 µl or 1 ml of 40% acrylamide mixed with 250 µl or 1 mL of 2% bis-acrylamide in 5 ml total volume to produce poly-acrylamide solutions for 1.5 and 14 kPA gels, respectively. Two hundred nm fluorescent beads (ThermoFisher) were mixed upon gel polymerization with ammonium persulfate and TEMED. Hydrogels were flattened using Sigmacote-treated (Sigma-Aldrich) 12 mm glass coverslips and turned upside down during polymerization to allow accumulation of fluorescent beads at their surface. Gels were activated with 0.5 mg/ml sulpho-SANPAH for 20 minutes under UV light and functionalized with 10 µg/ml fibronectin. Gel stiffness was checked by AFM. Live cells were stained with 0.5 µM SirActin probe (Cytoskeleton) or 2.5 µM CellMask (ThermoFisher Scientific). Imaging was performed after 60 min of cell adhesion and spreading with an inverted Apotome2 - Z1 Axio Observer microscope (Zeiss) mounted with a cMOS camera (Hamamatsu) and equipped with an incubator (Pecon) maintaining cells at 37°C and 5% CO_2_. Cells were imaged with a Plan Apochromat 40x water immersion objective. Multi-position z-stacks of images of beads and cells were acquired before detaching cells using SDS 20% (stressed state) and performing a second acquisition of the beads alone (relaxed state). Image analysis was done using an in-house Matlab (Mathworks) code. Bead images were filtered by means of a difference of Gaussian filter to enhance sphere-like structures and suppress noise. A binary mask of the cells was obtained by means of thresholding. Global rigid registration based on phase correlation was applied to correct for possible stage shifts between the stressed and relaxed images. Cell-induced displacements were calculated by means of Free Form Deformation-based non-rigid registration (FFD), as described in previous studies(*66*, *68*, *69*). Briefly, the Matlab orchestrated code calls the open-source image registration toolbox Elastix. Advanced Normalized Correlation was used as a registration metric and the quasi-Newton method for optimization. FFD uses a grid of control points that define a B-spline non-rigid transformation. Typically, a grid size between 8 and 16 pixels provides acceptable registration results. We used a grid size of 15 pixels for all the cases, which guaranteed that sufficient beads were present in between control points and provided successful registration metric results (around 99% for most of the cases)(*70*).

Tikhonov regularized Fourier Transform Traction Cytometry algorithm was applied assuming a homogeneous, isotropic and linear elastic half space. We used a previously proposed approach for which the amount of regularization does not scale with the Young’s modulus.

Choosing the amount of regularization (typically represented by the Greek letter λ) in studies that compare different conditions is non-trivial and no clear consensus has been reached in the field. To avoid biasing our conclusions we developed an automated method to obtain the value of λ that maximizes the following function:

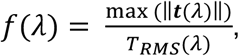

where ‖t(λ)‖ is the L2 norm of the traction vectors in every pixel in the field of view calculated with a given λ value, and T_RMS_ (λ) is the root mean square of ‖t(λ)‖. By finding the peak of f(λ) one obtains a λ that provides results with low level of background noise (low T_RMS_ (λ)) while preserving the higher traction values in the field of view (i.e., high max(‖T(λ)‖)), which are typically associated with the cell-induced tractions. Finally, cell tractions were recovered using the regularized inversion of the elasticity problem in Fourier domain. An Otsu thresholding algorithm was applied to the recovered tractions to obtain the *traction footprint* region(*66*). The metrics shown in Fig. 5, namely, average displacements (!𝒖_avg_), total force magnitude (‖𝒇_tot_‖) and average traction (!𝒕_avg_) were calculated within this region using the following formulas:

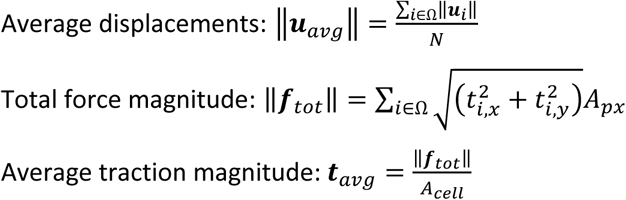

Where Ω is the traction footprint region, N is the number of pixels in Ω, 𝐭_4_ and 𝐮_4_ are the traction and displacement vectors, respectively, at spatial point i, and A_56_ is the area of the pixel.

### Stiffness-clamp experiments

Measurements for stiffness clamp were carried out as previously described(*39*, *71*). Briefly, experiments were conducted using a NanoWizard 4 BioAFM (JPK) associated with the CellHesion 200 module (JPK). Tipless cantilevers Arrow TL1 (NanoWorld) were coated with 40 µg/mL fibronectin for 20 min at room temperature. Cells were seeded in a 35mm µ-dish coated with fibronectin for 2 min. The cantilever was placed in contact with individual cells, and measurements were started after 150 s and ended within 30 min after capture on the cantilever.

### Atomic force microscopy and correlative atomic force and fluorescence microscopy

Topography of CDM was determined using a scanning force microscope NanoWizard 4 BioAFM (JPK) mounted on a Z1 Axio Observer epifluorescence microscope equipped with an Apotome2 (Zeiss). Measurements were performed using a conical shape probe with typical tip radius of curvature < 10 nm mounted on a gold coated cantilever (0.03-0.09 N/m) (Biolever Mini, Olympus). Spring constant was determined upon calibration by the thermal noise method. Quantitative imaging (QI) (JPK, Berlin, Germany) was conducted in PBS. Typically, an AFM map of 20 x 20 μm corresponding to 256x256 pixels was acquired at an appropriate scan speed (100 to 200 µm/s) and the lowest setpoint possible (typically 0.1 nN) to prevent damage to the sample. AFM image processing and analysis were performed using JPK data processing software and ImageJ. Second order flattening was applied to AFM topography images to remove tilt and bow. Median filter and Gaussian convolution were also applied when necessary. Matrix coverage was assessed as the proportion of pixels higher than 50 nm. Mean height was calculated for 75% of the highest fibers. Acquisitions and processing for correlation with fluorescence were performed based on the optical calibration module (JPK). Quantification of fluorescence content was achieved as described (Fig. S3) by thresholding both AFM and fluorescence images and calculating integrated density of pixelized images.

### Measurement of barrier properties

Endothelial barrier properties were measured using the impedance-based xCELLigence cell analyzer (RTCA system, Agilent) at 10 kHz according to the manufacturer’s recommendations. Briefly, endothelial cells were seeded in complete ECGM2 medium at confluency in 16-well E-plates. Preliminary experiments included assessment of cell density for formation of a monolayer early after seeding, without overseeding that would result in overlapping cells. Barrier properties were followed over time long after the plateau was reached. Basal barrier properties were measured in three independent experiments in quadruplicate wells each at 72 h. In some experiment cells were then treated or not with 2 U/ml thrombin (Merck Millipore) in duplicate wells to assess cell response to regulator of permeability. Cell index was thus normalized to the plateau value before addition of thrombin and live measurements were performed over the next 4h.

### Protein extraction and immunoblotting

Endothelial cells were lysed with Laemmli buffer containing 4% SDS and 50 mM DTT before separation by SDS-PAGE. Proteins were transferred to PVDF membranes before incubation with the appropriate antibodies in 25 mM Tris-HCl, pH 7.4, 0.1% Tween 20, and 5% milk. Immunological detection was performed with horseradish peroxidase (HRP) conjugated secondary antibodies, using ECL (Life Technologies) as a substrate. Pixel intensity was measured using imageJ software.

### 3D microgrooved substrates experiments

Topographic microgrooved substrates were prepared as already described(*61*). Briefly, an original microstructured silicon wafer is designed using classical photolithography procedures by UV illumination. The final microstructured culture coverslip is generated from a PDMS mold made from the original wafer. The microstructures consist of parallel arrays of rectangular grooves, with a fixed depth of 1 µm and with a width and spacing of 2 or 5 µm. Each coverslip included an unpatterned area of PDMS substrate that was used as control. The coverslips were plasma-cleaned for 30 seconds before directly seeding confluent cells in ECGM2. Cell monolayers were fixed with 4% PFA after 24, 48 or 72 h and immunostained as described. For image analysis, the cell outlines were segmented from VE-cadherin staining using the Tissue Analyzer plugin (ImageJ). Cell shape parameters were then extracted from the cell outlines using custom-written Matlab (MathWorks) scripts(*61*). Absolute orientation describes the cell orientation angle between 0° (in the direction of the grooves) and 90° (perpendicular to the grooves). Circularity describes the cell aspect ratio when fitted with an ellipse (ratio of minor to major axis).

### Statistical Analysis

Experiments were performed at least three times and included independent culture duplicates unless otherwise indicated in figure captions. Data are presented as mean ± SD, and statistical significance was determined using the specific tests indicated in the figure captions. In all cases in which a test for normally distributed data was used, normal distribution was confirmed using D’Agostino–Pearson or Shapiro-Wilk tests. All calculations were performed using Prism 8.0 (GraphPad Software). *p* values are described in the figure captions or stated on the figure itself.

## ACKNOWLEDGEMENTS

We thank Simone Bovio (Platim, ENS Lyon) and Pierre-Henri Puech (LAI Lab, Marseille) for helpful discussion concerning AFM experimental design and correlative microscopy, as well as Torsten Mueller (JPK BioAFM Center, Brucker Nano Gmb) for constant support. We thank Anaïs Monet (MBI, National University of Singapore) for providing the python patching script. HUVEC were prepared from umbilical cords obtained through AP-HP, Hôpital Saint-Louis, Unité de Thérapie Cellulaire, CRB-Banque de Sang de Cordon, Paris, France – N° d’autorisation : AC-2016-2759. We thank Emmanuel Pauthe (ERRMECe, Université de Cergy) for the gift of human plasma fibronectin and Christophe Helary (LCMCP, Sorbonne University, Paris) for the gift of rat tail collagen I. This work was supported by funding from an ERPT grant from Fondation Leducq. MFM was supported by Fondation pour la Recherche Médicale, YA by Ligue Nationale contre le Cancer, CUD and NBJ by Sorbonne Université and fondation ARC Jeunes Chercheurs (NBJ), CL by Fondation Lefoulon-Delalande and Fondation Bettencourt-Schueller, and AIB by an endowment in cardiovascular bioengineering from the AXA Research Fund. JBF is supported by an FWO postdoctoral fellowship 1259223N. AS is supported by the FWO SB grant 1S68818N. HVO acknowledges funding from FWO G0C2422N and KU Leuven internal funding C14/17/111.

## AUTHOR CONTRIBUTIONS

Conceived, designed and directed the study: MFM, LM Performed experiments: MFM, NBJ, CL, YA, CUD, CAR, SM, LM Performed the stiffness-clamp experiments: CG Contributed with AFM acquisitions and image analyses: TP, PM Analyzed data: MFM, NBJ, CL, JBF, AS Provided essential protocols and contributed to design of the study: AIB, CM, HVO, SG Contributed to manuscript preparation: CL, SG, CM Wrote the manuscript: MFM, LM Administration and funding of the study: LM, SG

## ABBREVIATIONS

AFM: atomic force microscopy
BM: basement membrane
CDM: cell-derived matrix
ECM: extracellular matrix
ERK: extracellular signal regulated kinase
FAK: focal adhesion kinase
HUVEC: human umbilical vein endothelial cell
LOXL2: lysyl oxidase like-2
VEGF: vascular endothelial growth factor

## Notes

### Competing Interest Statement

The authors have declared no competing interest.

