## Supplemental figures for "Adaptation of endothelial cells to microenvironment topographical cues through lysyl oxidase like-2-mediated basement membrane scaffolding"

**SUPPLEMENTARY FIGURE 1**

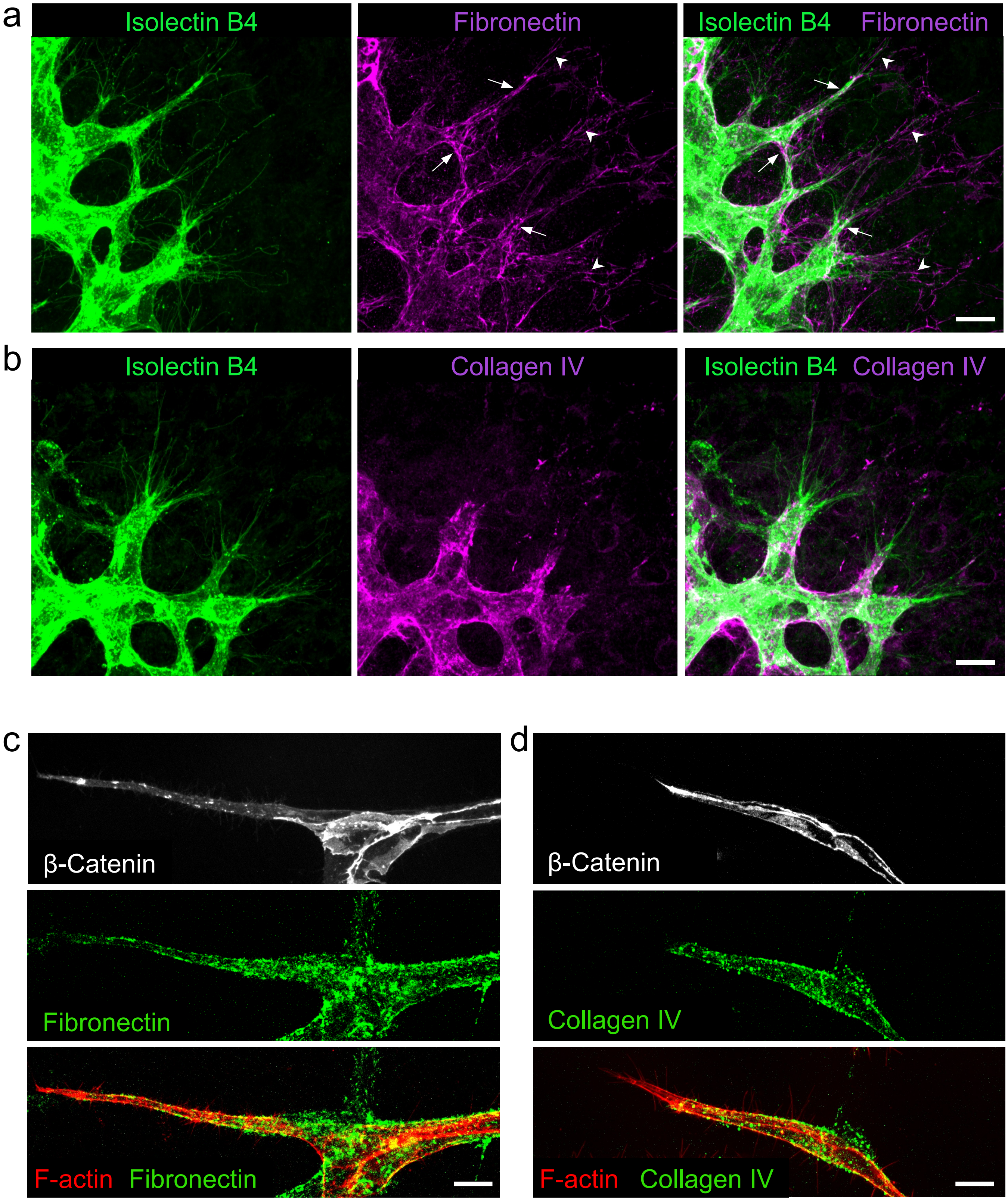


**Fig. S1. Angiogenesis-associated basement membrane deposition**
**a-b**: Immunostaining of fibronectin (**a**) and collagen IV (**b**) (magenta) in P5 mouse retina. Endothelial cells were detected using Griffoniae simplicifolia Isolectin B4 (green). White arrowheads point to fibronectin deposits in the avascular microenvironment and white arrows to endothelial fibronectin in the basement membrane. Scale bar: 10 µm. **c-d**: Endothelial cells were seeded in collagen hydrogel and cultured for 72 h to generate *in vitro* capillary-like structures. Immunostaining of fibronectin (**c**) and collagen IV (**d**) (green) in *in vitro* capillary-like structures. Endothelial cell junctions were immunostained for β-Catenin (white) and F-actin was detected using phalloidin (red). Scale bar: 40 µm. Confocal stacks of images were acquired and the projections are illustrated (**a-d**).

**SUPPLEMENTARY FIGURE 2**

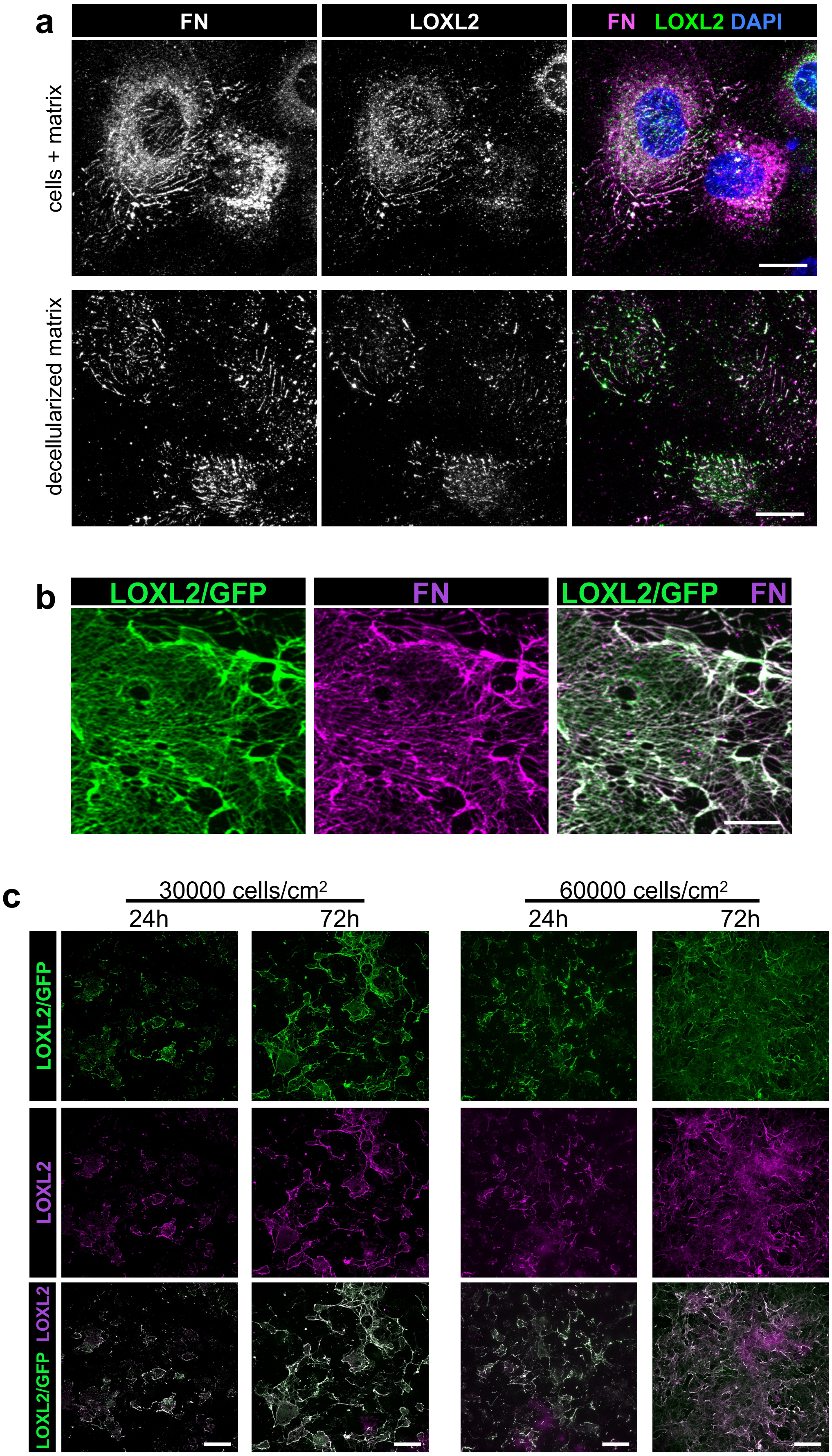


**Supplementary figure 2. Characterization of cell-derived matrices.** **(a)** Immunostaining of fibronectin (white in left column and magenta) and LOXL2 (white in central column and green) in endothelial cell culture (top row) and in CDM (bottom row) prepared 6 h after seeding cells. Nuclei were stained with DAPI (blue). Scale bar: 20 µm. **(b)** Immunostaining of fibronectin (magenta) in CDM prepared 72 h after seeding endothelial cells expressing LOXL2/GFP. Scale bar: 20 µm. **(c)** Immunostaining of LOXL2 (magenta) in CDM prepared 24 or 72 h after seeding endothelial cells expressing LOXL2/GFP (green) below (30 000 cells/cm$^2$) or at (60 000 cells/cm$^2$) confluency. Scale bar: 100 µm.

**SUPPLEMENTARY FIGURE 3**

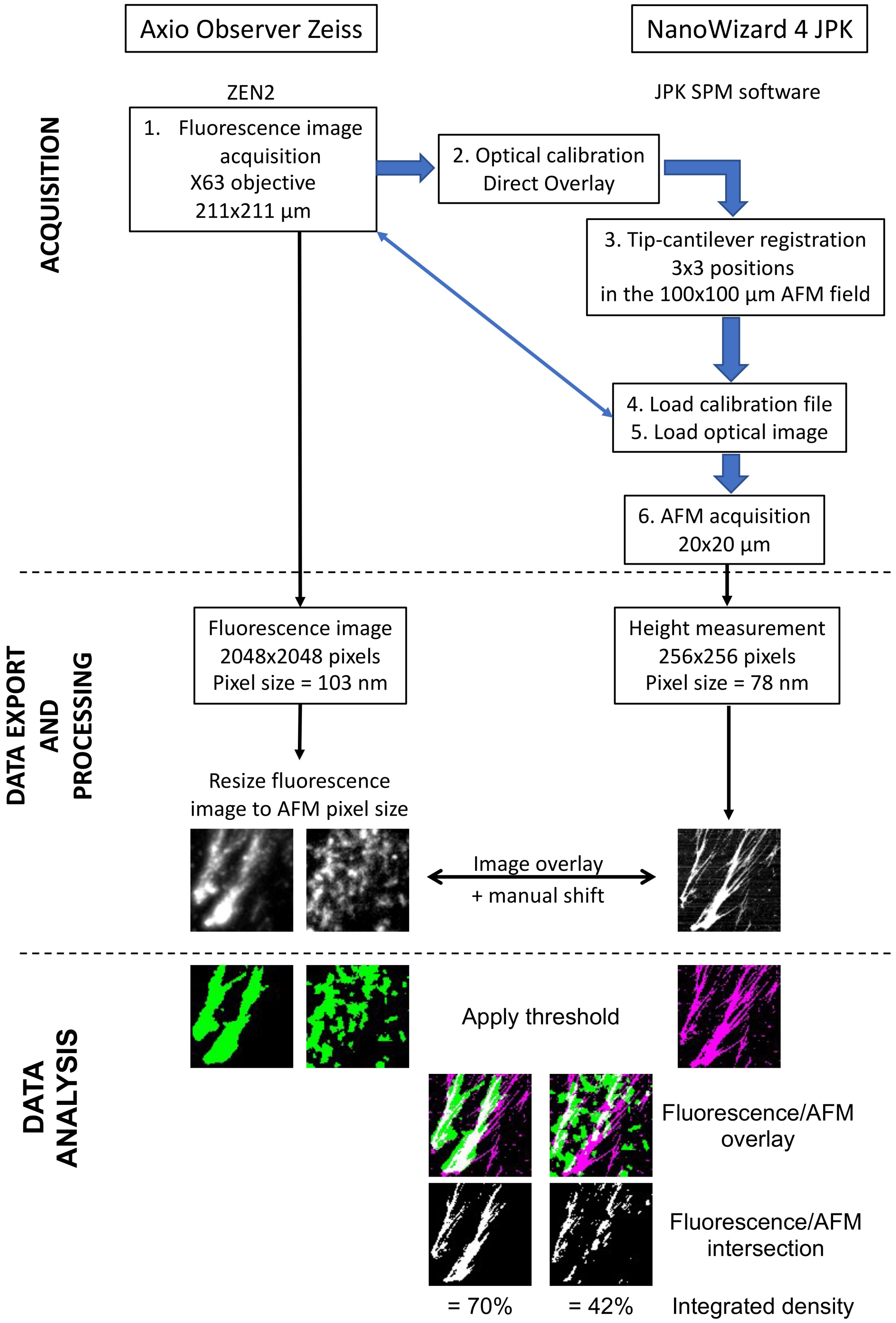


**Supplementary figure 3. Correlative AFM and fluorescence microscopy.** Acquisitions were performed using the Zen2 software to control the Zeiss Observer fluorescence microscope and the SPM JPK software to control the Nanowizard 4 AFM microscope. Briefly, a fluorescence image was acquired and imported in the JPK software to perform the AFM acquisition. Image overlay was achieved by matching pixel size of the two images before data export and included a manual translation step using a recognizable pattern. To quantify the correlation rate between the fluorescence signal and the AFM-detected structures, fluorescence images were cropped, resized to similar pixel dimension and manually thresholded for all channels using ImageJ software before calculation of integrated density. Proportion of pixels corresponding to AFM detected structures and containing fluorescent signal was then calculated as the ratio of integrated density.

**SUPPLEMENTARY FIGURE 4**

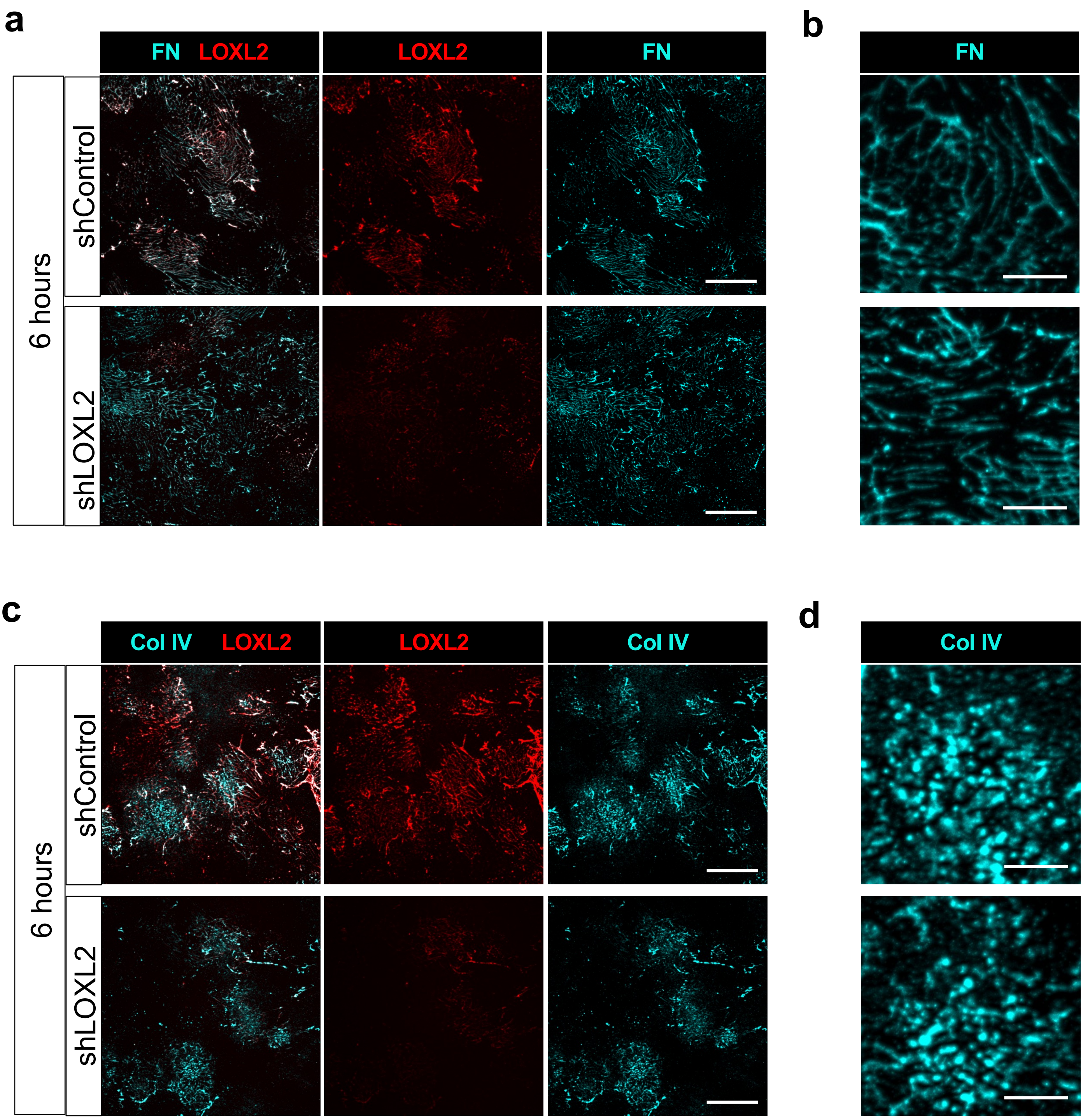


**Supplementary figure 4. LOXL2 does not affect deposition of basement membrane components at early timepoints. (a-b)** Immunostaining of fibronectin **(a and b)** or collagen IV **(c and d)** (cyan) and LOXL2 (red) in CDM generated by control (shControl) or LOXL2-depleted (shLOXL2) endothelial cells maintained for 6 h. Scale bar: 50 (a and c) and 10 (b and d) µm.

### SUPPLEMENTARY FIGURE 5

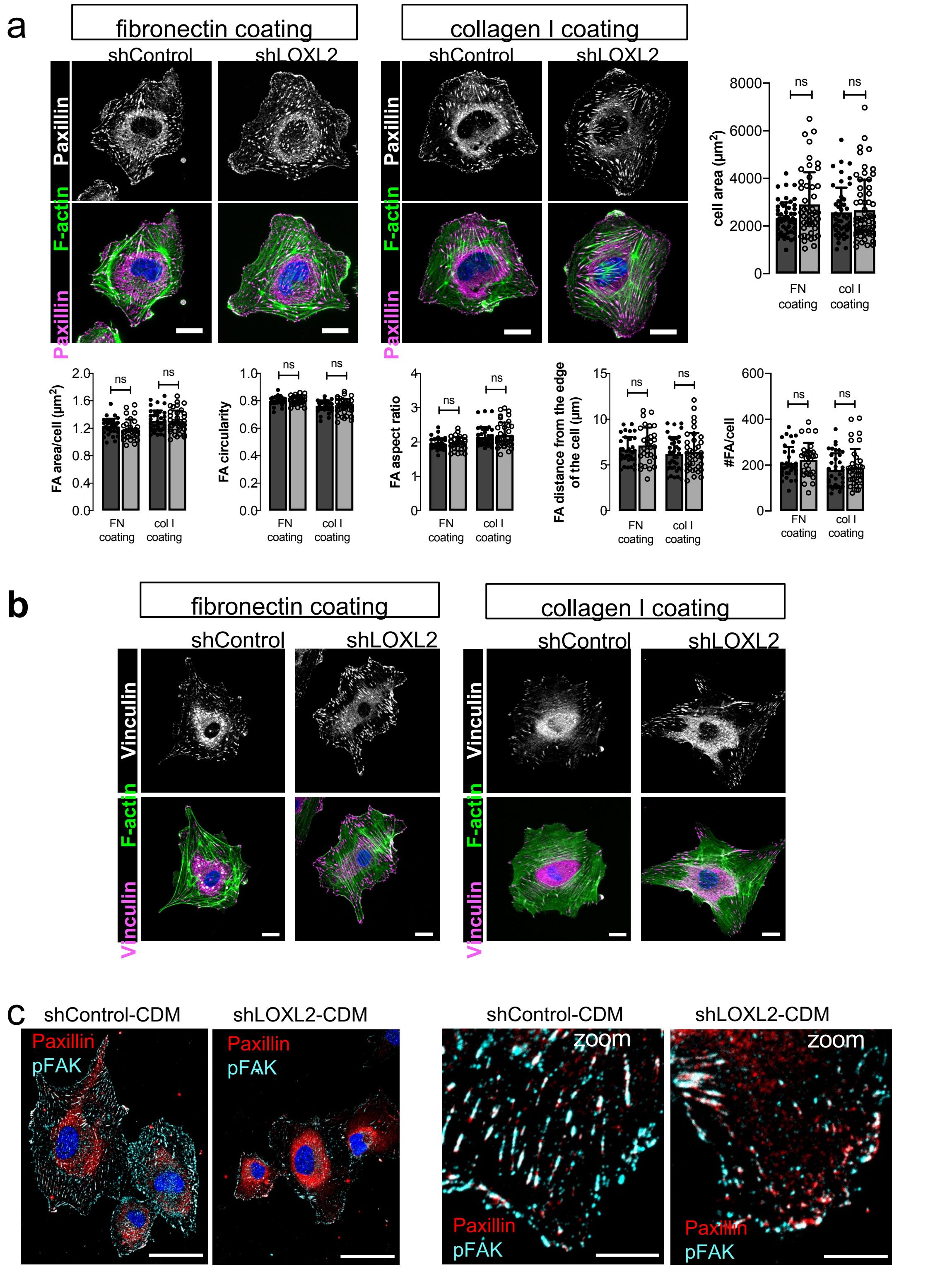


**Supplementary figure 5. LOXL2 depletion *per se* does not alter the adhesion and contractility capacities of endothelial cells.**

**(a)** Paxillin was immunostained (top row - magenta) in control (shControl) and LOXL2-depleted (shLOXL2) endothelial cells plated for 1 h on fibronectin or collagen I surface coating. F-actin was detected with phalloidin-Alexa Fluor 488 (green). Nuclei were stained with DAPI (blue). Scale bar: 10 µm. Surface area of individual cells and morphometric parameters of focal adhesions (mean area, circularity, aspect ratio, distance from the edge of the cell and number of focal adhesions per cell) were calculated in approximately 60 cells from two independent experiments. Statistical analysis was performed using unpaired t- tests. **(b)** Vinculin was immunostained (top row - magenta) in control (shControl) and LOXL2-depleted (shLOXL2) endothelial cells plated for 1 h on fibronectin or collagen I surface coating. F-actin was detected with phalloidin-Alexa Fluor 488 (green). Nuclei were stained with DAPI (blue). Scale bar: 10 µm. **(c)** Immunostaining of paxillin (red) or pY397-FAK (pFAK - cyan) in control cells 1 h after plating on CDM prepared from control or LOXL2-depleted endothelial cells cultivated for 72 h. Scale bars: 50 (left panel) and 10 (right panel) µm.

**SUPPLEMENTARY FIGURE 6**

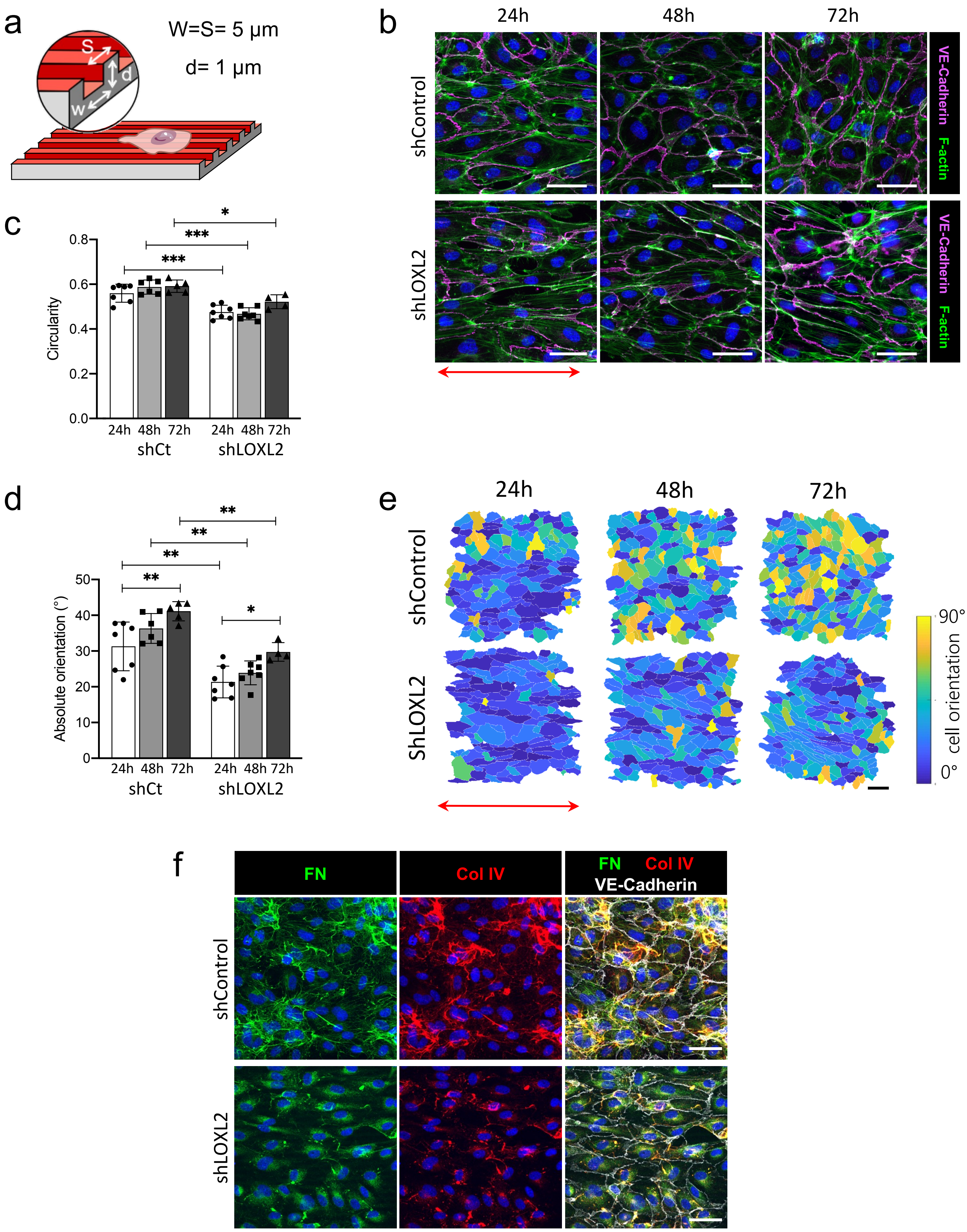


**Supplementary figure 6. Defective basement membrane scaffolding affects cell response to topography (a)** Schematic of the microgrooved culture substrate. **(b-e)** Immunostaining of VE-cadherin (magenta) and staining of F-actin with phalloidin-Alexa Fluor 488 (green) in control (shControl) or LOXL2-depleted (shLOXL2) endothelial cells 24, 48 or 72 h after seeding at confluency on microgrooved coverslips **(b).** Scale bar: 50 µm. Cell shape **(c)** and orientation **(d-e)** were measured after segmentation and image analysis in 3 independent experiments. Scale bar: 100 µm. Double-arrow indicates the orientation of the microgrooves **(f)** Immunostaining of fibronectin (green), collagen IV (red) and VE-cadherin (white) in cells cultured for 72 h on microgrooved substrate. Scale bar: 50 µm. Nuclei were detected with DAPI (blue).

**SUPPLEMENTARY FIGURE 7**

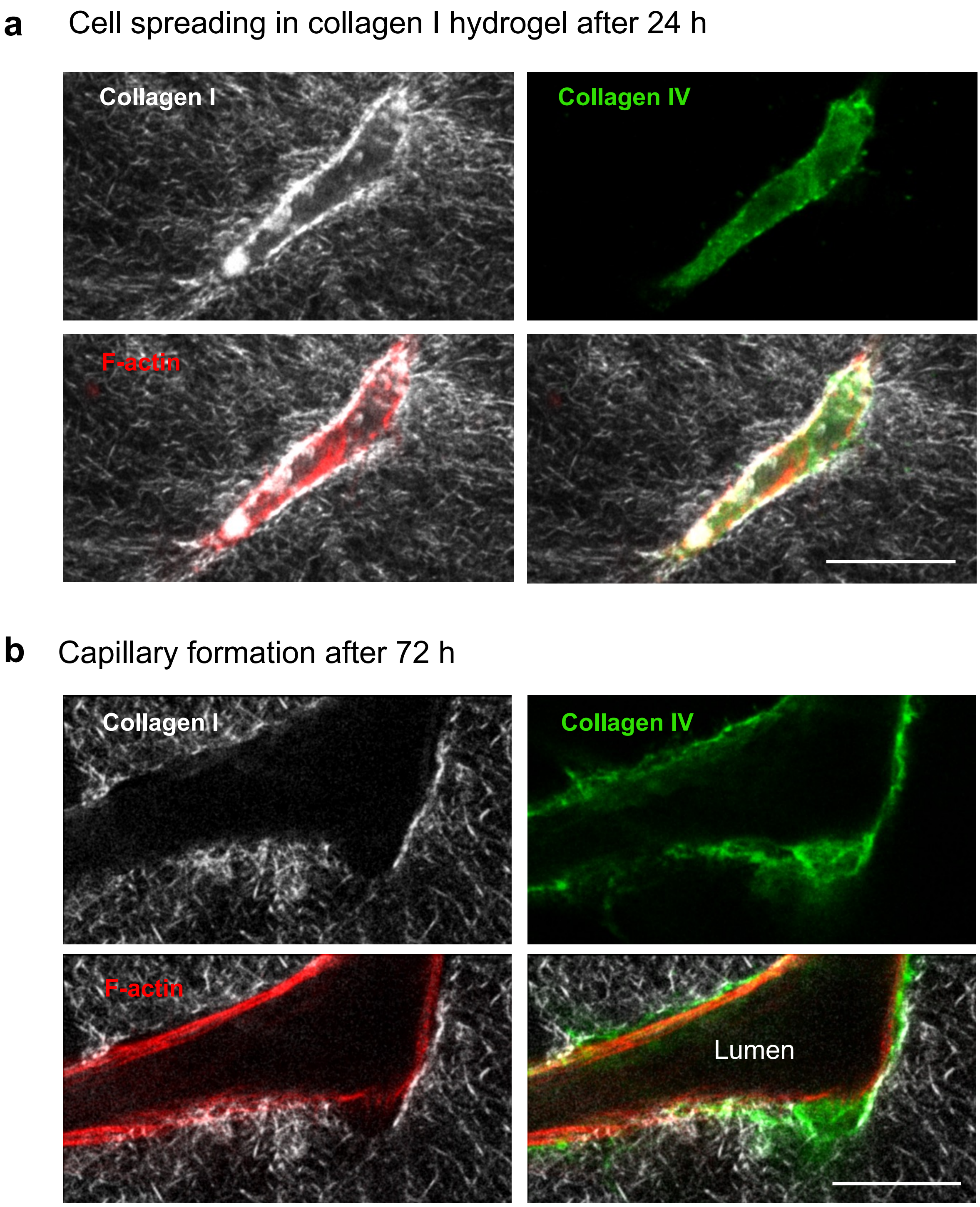


**Supplementary figure 7. ECM remodelling at the surface of endothelial cell during capillary formation.** Detection of fibrillar collagen using SHG in 2-P microscopy (white) and immunostaining of collagen IV were performed to assess extracellular matrix remodeling associated to capillary formation 24 **(a)** and 72 **(b)** h after seeding in collagen I hydrogels. F-actin was labelled with phalloidin (red). Single confocal planes are shown. Scale bar: 25 µm.

**SUPPLEMENTARY FIGURE 8**

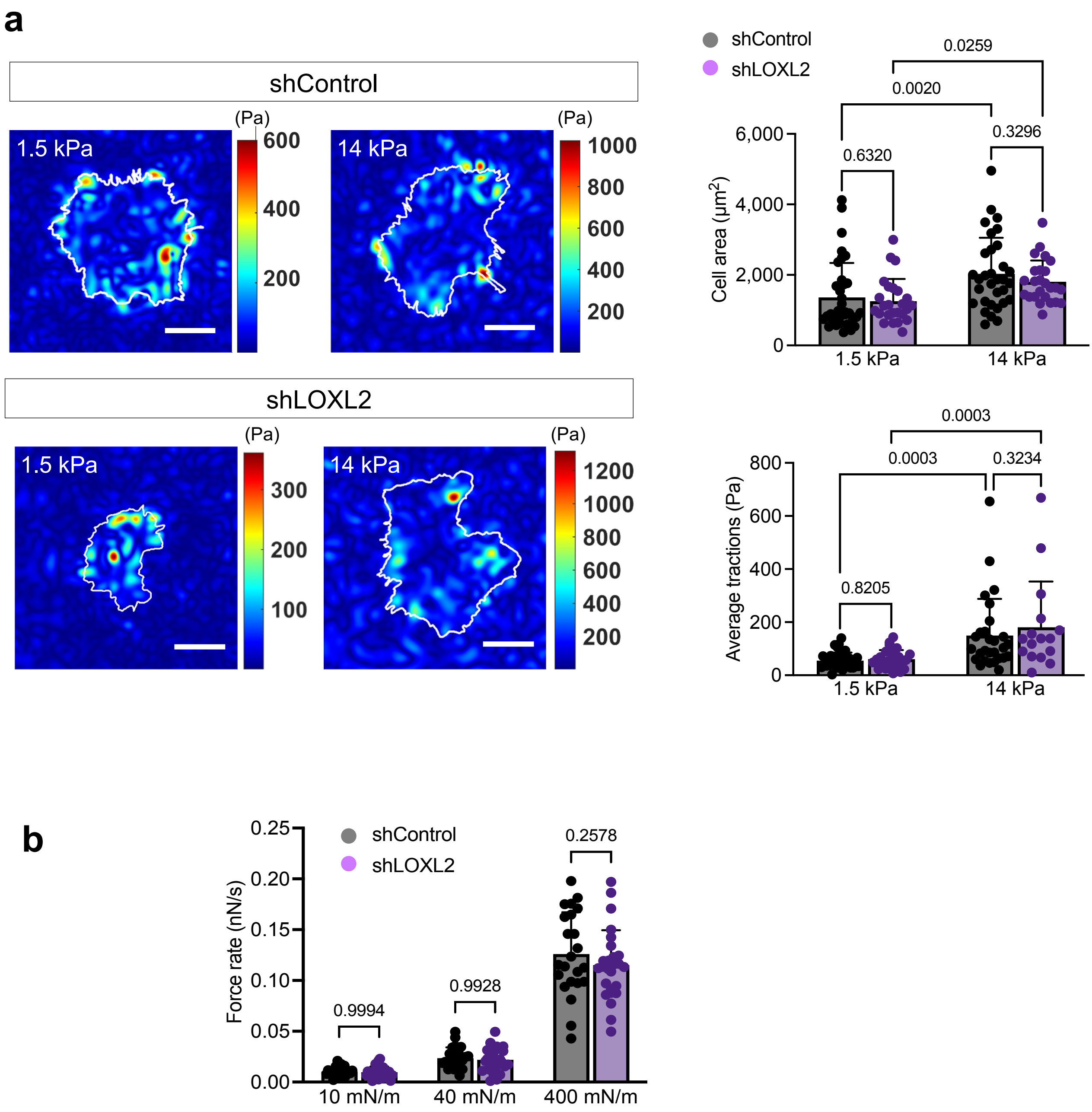


**Supplementary figure 7. Basement membrane deposition and 3D vascular morphogenesis**

**(a)** Traction force microscopy measurements were performed 60 min after cell seeding on 1.5 and 14 kPa fibronectin-coated polyacrylamide hydrogels. Heat maps of average tractions (Pa) are shown. Scale bar: 20 µm. Cell area (µm²) and average tractions (Pa) were calculated in approximately 30 cells from 4 independent experiments. Statistical analysis was performed using two-way ANOVAs. **(b)** Endothelial cells were captured on an AFM cantilever coated with fibronectin and immobilized on a fibronectin coating for measurement of their mean force rate in an AFM-stiffness clamp set-up. Results are presented as the mean of approximately 25 cells from 3 independent experiments. Statistical analysis was done using two-way ANOVA.

**SUPPLEMENTARY FIGURE 9**

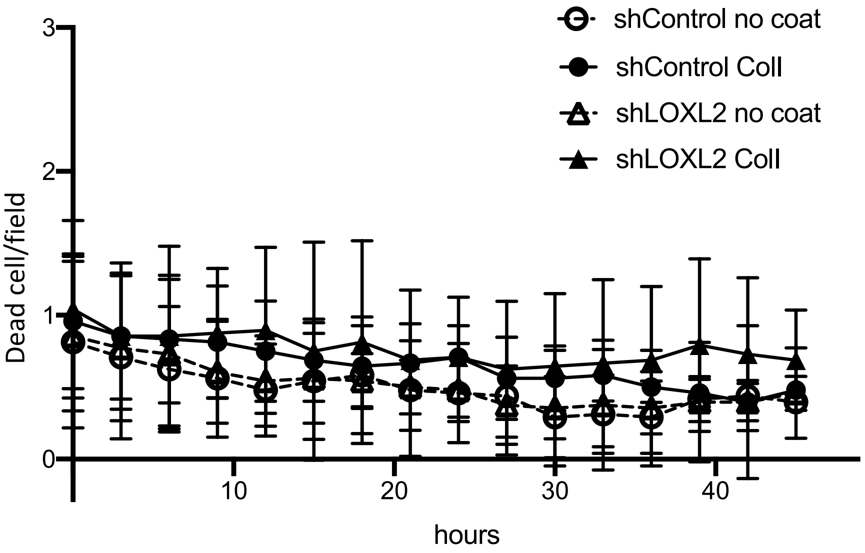


**Supplementary figure 9. LOXL2 does not impact cell death in 2D cell culture.** Control (circle) and LOXL2-depleted (triangle) endothelial cells were seeded on tissue culture plastic (open symbols) or collagen I coating (closed symbols). Cells were cultured in the presence of Cytotox Green dye (Sartorius) to detect dead cells over 48 h in a Zoom Incucyte cell imager (Essen Bioscience, Sartorius). Dead cells were counted using proprietary software (Essen Bioscience, Sartorius).
